# Molecular basis of differential adventitious rooting competence in poplar genotypes

**DOI:** 10.1101/2021.09.14.460203

**Authors:** Alok Ranjan, Irene Perrone, Sanaria Alallaq, Rajesh Singh, Adeline Rigal, Federica Brunoni, Walter Chitarra, Frederic Guinet, Annegret Kohler, Francis Martin, Nathaniel Street, Rishikesh Bhalerao, Valérie Legué, Catherine Bellini

## Abstract

- Recalcitrant adventitious root (AR) development is a major hurdle in propagating commercially important woody plants. Although significant progress has been made to identify genes involved in subsequent steps of AR development, the molecular basis of differences in apparent recalcitrance to form AR between easy-to-root and difficult-to-root genotypes remains unknown.
- To address this, we generated cambium tissue-specific transcriptomic data from stem cuttings of hybrid aspen, T89 (difficult-to-root) and hybrid poplar OP42 (easy-to-root) and used transgenic approaches to verify the role of several transcription factors (TF) in the control of adventitious rooting.
- Increased peroxidase activity is positively correlated with better rooting. We found differentially expressed genes encoding Reactive Oxygen Species (ROS) scavenging proteins to be enriched in OP42 compared to T89. A higher number of differentially expressed TF in OP42 compared to T89 cambium cells was revealed by a more intense transcriptional reprograming in the former. *PtMYC2*, a potential negative regulator, was less expressed in OP42 compared to T89. Using transgenic approaches, we have demonstrated that *PttARF17.1* and *PttMYC2.1* negatively regulate adventitious rooting.
- Our results provide insights into the molecular basis of genotypic differences in AR and implicate differential expression of the master regulator MYC2 as a critical player in this process.

## Introduction

In the 1990s, only 3% of the world’s forested land was as plantations. However, despite this small percentage, it still provided more than one third of total industrial wood production (Kirilenko and Sedjo, 2007). The shift of production from natural forests to plantations is projected to accelerate and is expected to rise to 75% in the 2050s (Kirilenko and Sedjo, 2007). Operating plantations is expensive and requires high productivity per hectare, which in turn requires good quality, i.e., genetically improved planting stock. Many forest companies are therefore currently considering clonal propagation in addition to, or in conjunction with, their breeding programmes. This aims to propagate elite genotypes from available genetic diversity and maximise the productivity of selected high-value hybrid clones (Bozzano *et al*., 2014). The genus *Populus* comprises about 30 species; its wood forms an abundant and renewable source of biomaterials and bioenergy (Ragauskas *et al*., 2006). The propagation of poplar species depends primarily on AR formation from detached stem cuttings (Dickmann, 2006) but one major constraint for vegetative propagation of some economically important elite genotypes is incompetence or rapid loss of capacity in forming AR (Bellini *et al*., 2014; Brunoni *et al*., 2019; Bannoud and Bellini 2021). AR development is a complex, heritable trait controlled by many endogenous regulatory factors and much influenced by environmental factors (Bellini *et al*., 2014; Bannoud and Bellini 2021). The rooting capacity of cuttings varies among individuals within species, populations, or even clones (Abarca and Díaz-Sala, 2009a; Abarca and Díaz-Sala, 2009b). Few studies have reported the genetic variability of AR development of *Populus* hardwood cuttings. Zhang *et al*. (2009), reported quantitative trait loci (QTL) that control two AR growth parameters in a full-sib family of 93 hybrids derived from an interspecific cross between two *Populus* species, *P. deltoides* and *P. euramericana*, which are defined as difficult-to-root and easy-to-root, respectively. They showed that the maximum root length and the total number of AR were correlated and under strong genetic control, which supports earlier genetic QTL analysis performed on forest trees (reviewed in Geiss *et al*., 2009). Several studies focusing on AR development in poplar have identified a number of genes involved in its regulation (Ramirez-Carvajal *et al*., 2009; Rigal *et al*., 2012; Trupiano *et al*., 2013; Wuddineh *et al*., 2015; Li *et al*., 2018; Liu *et al*., 2020; Wang *et al*., 2020; Wei *et al*., 2020; Xu *et al*., 2021; Xu *et al*., 2015; Yordanov *et al*., 2017; Yue et al., 2020; Zhang *et al*., 2020) including large-scale data analyses identifying regulators (Ribeiro *et al*., 2016; Zhang *et al*., 2019) and pharmacological assays of physiological regulators (Gou *et al*., 2010; Mauriat *et al*., 2014; Zhang *et al*., 2019). All these studies resulted in a substantial increase in our understanding of the molecular mechanisms that control successive steps of AR development, but the molecular basis of recalcitrance to form AR between easy-to-root and difficult-to-root genotypes remains unknown. To address this question, we compared the transcriptome of cambium cells obtained immediately after cutting and 24 h later by Laser Capture Microdissection (LCM) from *P. trichocarpa* × *P. maximowiczii* (clone OP42) which we defined as ‘easy-to-root from woody stem cuttings’, and the hybrid aspen *P. tremula* × *P. tremuloides* (clone T89) which we defined as ‘difficult-to-root from woody stem cuttings’. OP42 is one of the poplar clones used most widely, both in Northern Europe and worldwide (Taeroe *et al*., 2015). It can easily be propagated from dormant stem cuttings. By contrast, the hybrid aspen T89 cannot be propagated *via* dormant stem cuttings but can be easily propagated *in vitro* and is very amenable to genetic transformation (Nilsson *et al*., 1992). The analysis of the transcriptomic Dataset showed there to be more differentially expressed transcription factors (TF) in OP42 than in T89. We identified several TF that could explain differences in aptitude to produce adventitious roots. We show that the up-regulation of the jasmonate (JA) signalling pathway in the cambium of T89 could be one cause of the failure to produce adventitious roots.

## Materials and Methods

### Plant growth conditions and rooting assays

The hybrid aspen (***P. tremula** L*. × ***P. tremuloides** Michx*), clone **T89, and** the hybrid poplar (*P.trichocarpa*× *P. **maximowiczii***) clone **OP42, were** propagated *in vitro* for four weeks as described in Karlberg *et al*., (2011) (Methods S1; Fig. S1a). For *in vitro* rooting assays, 3 cm cuttings of T89 and OP42 plantlets were collected and transferred to fresh sterile medium (Methods S1; **Fig.** S1b, d). The number of AR was scored from day five after cutting until day 14. Three replicates of 15 stem cuttings each were analysed. For the jasmonic acid treatment, cuttings from four-week-old *in vitro* T89 and OP42 plantlets were transferred to fresh sterile medium with or without methyl jasmonate (MeJA) at 5 μM, 10 μM, or 20 μM.

For the rooting assay in hydroponic conditions, 20 cm long stem cuttings taken from the third internode below the shoot apex from three-month-old T89 and OP42 plants grown in the greenhouse, were transferred to hydroponic conditions (Methods S1; **Fig.** S1c,e;).

### Histological analysis of stem cuttings *in vitro*

5 mm stem fragments were taken at the base of cuttings four or five days after cutting. Samples were fixed and prepared for sectioning as described in Methods S2. 10 μm sections were obtained with a rotary microtome (https://www.zeiss.com/) and stained with safranin (Sigma-Aldrich, https://www.sigmaaldrich.com/) and alcian blue (Sigma-Aldrich, https://www.sigmaaldrich.com/) in a ratio of 1:2; using methods from Hamann *et al*., (2011).

### Tissue preparation before LCM

#### Sampling, fixation and cryoprotection steps

The basal 5 mm stem segments from T89 and OP42 cuttings were harvested immediately after excision from greenhouse grown plants (Time T0) and after 24 h of hydroponic culture (Time T1) (Fig. S2a-c). For each genotype, at each time point, three biological replicates were collected (12 stem segments in total = 3 biological replicates × 2 genotypes × 2 time points). Immediately after sampling, the stem pieces were split in half longitudinally, subjected to fixation and cryoprotection steps before the laser microdissection according to the protocol described at https://schnablelab.plantgenomics.iastate.edu/resources/protocols/, slightly modified as described in Methods S3.

#### Cryosectioning

Samples were fixed with Tissue-Tek® Optimal Cutting Temperature (O.C.T.) compound onto a specimen stage directly in the cryo chamber. Stem segments were mounted to allow cryosectioning, and cambium collection from tangential cryosections (Fig. S2d) in order to avoid embedding and the presence of O.C.T. compound on membrane slides. Sections of 25 μm were transferred onto membrane slides. Three progressive dehydration steps in ethanol were applied. After ethanol removal, sections were air-dried before microdissection (Methods S3).

### Laser capture microdissection, RNA extraction, and RNA Sequencing

LCM was performed with a PALM Robot-Microbeam system (Zeiss MicroImaging, Munich, Germany). Cambium microdissected cells were catapulted into the adhesive caps of 500 μl tubes (Zeiss) (Fig. S2e-k). Total RNA was isolated using the PicoPure RNA Isolation Kit (Thermo Fisher Scientific, https://www.fishersci.se/se/en/home.html). Quality and quantity of RNA samples were assessed using the Bio-Rad Experion analyser and Experion RNA high-sense analysis kit (Bio-Rad). Total RNA from each biological replicate was amplified using the MessageAmp II aRNA amplification kit (Ambion, Austin, TX, U.S.A.). Amplified RNA profiles were verified using the Experion analyser and Experion RNA standard-sense analysis kit (Bio-Rad). In total, twelve cDNA paired-end libraries were generated using the mRNA-Seq assay for transcriptome sequencing on an Illumina HiSeq™ 2000 platform at Beijing Genome Institute (BGI, China), but only eleven were sequenced as one T89 (T1) sample failed the quality check.

### Pre-processing of RNA-Seq data

The data pre-processing was performed as described in: http://www.epigenesys.eu/en/protocols/bio-informatics/1283-guidelines-for-rna-seq-data and detailed in Methods S4.

### Differential gene expression analysis

Statistical analysis of single-gene differential expression between conditions was performed in R (v3.4.0; Team, 2018) using the Bioconductor (v3.5; Gentleman *et al*., 2004) DESeq2 package (v1.16.1; Love *et al*., 2014). FDR adjusted p-values were used to assess significance; a common threshold of 1% was used throughout. For the data quality assessment and visualisation, the read counts were normalised using a variance stabilising transformation (vst) as implemented in DESeq2. The biological relevance of the data, such as similarity of biological replicates (Fig. S3a,b) and other visualisations (e.g., heat maps), were obtained using custom R scripts, available at https://github.com/nicolasDelhomme/poplar cambium.

Dendrograms and heat maps were generated using the function heatmap.2 from the gplots R library. Heat maps of differentially expressed genes (DEG) (DE cut-offs of FDR ≤ 0.01 and |LFC| ≥ 0.5), were generated using the function heatmap.2 from the gplots R library. The 17,997 genes, which were detected in all biological replicates, were used for further analysis. Genes which were expressed only in one or two biological replicates for each genotype, but which were significantly differently expressed between T89 and OP42, were analysed separately. The gene expression mean values are listed in Dataset S3 (sheet 6).

### Gene ontology analysis

The REVIGO web server (http://revigo.irb.hr/) was used to summarise GO terms from differentially expressed genes (Supek *et al*., 2011). The GO terms with a false discovery rate (FDR; e-value corrected for list size) of ≤ 0.05 were submitted to the REVIGO tool, and the ‘small allowed similarity’ setting was selected to obtain a compact output of enriched GO terms. The overall significance of enriched processes was expressed as the sum of 100 × -log_10_ (FDR) for each enriched GO term counted within that process. This technique was adapted from the method used to visualise enriched GO terms as a percentage of the total enriched terms in the TreeMap function of the REVIGO web server.

### Transcription factors and digital differential gene expression analysis

The gene list of *P. trichocarpa* transcription factors was downloaded from the plant transcription factor database v4.0 (http://planttfdb.gao-lab.org/).

### Identification of poplar homologues of Arabidopsis *ARFs* and *MYC* transcription factors

Full-length amino acid sequences of the selected poplar and Arabidopsis *AUXIN RESPONSE FACTOR* (*ARF*) genes were subjected to phylogenetic analysis as described in Methods S5. The most closely related orthologues were chosen for the study (Fig. S4a). We used poplar *ARF* gene names according to the nomenclature in PopGenIE. Corresponding gene names are as follows: *PtrARF6.1*; Potri.005G207700, *PtrARF6.2*; Potri.002G055000, *PtrARF6.3*; Potri.001G358500, *PtrARF6.4*; Potri.011G091900, *PtrARF8.1*; Potri.004G078200, *PtrARF8.2*; Potri.017G141000, *PtrARF17.1*; Potri.005G171300 and *PtrARF17.2*; Potri.002G089900. Similarly, the poplar homologues of Arabidopsis *AtMYC2.1* were analysed; their corresponding gene names are as follows: *PtrMYC2.1*; Potri.003G092200, *PtrMYC2.2*; Potri.001G142200, *PtrMYC2.3;* Potri.002G176900, *PtrMYC2.4*; Potri.001G083500, *PtrMYC2.5;*Potri.003G147300 and *PtrMYC2.6*; Potri.014G103700.

### Generation of transgenic hybrid aspen plants

To amplify the candidate genes, cDNA was synthesised starting from total RNA extracted from hybrid aspen T89 (*P. tremula* × *P. tremoloides*) leaves and DNAse treated. As it is not possible to distinguish the sequence of *P. tremula* from that of *P. tremuloides*, the genes are referred to as *PttARF6.4, PttARF8.2, PttARF17.2* and *PttMYC2.1* and the corresponding primers used for amplification of the coding sequence are listed in Table S1.

Transgenic T89 plants over-expressing *PttARF6.4, PttARF8.2* or *PttMYC2.1* or down-regulated for the expression of *PttARF6.4, PttARF8.2* and *PttARF17.2* were produced as described in the Methods S5. The relative expression levels of *PttARF6.1/2, PttARF6.3/4, PttARF8.1/2, PttARF17.1/2* and *PttMYC2.1* in the respective transgenic lines were further quantified by qPCR.

### Quantitative Real-Time PCR analysis

To check the over-expression or the down-regulation of the selected genes in the transgenic lines, five 5 mm stem pieces were taken at the base of cuttings from T89 (3 biological replicates) and transgenic lines (3 biological replicates for each line) at the time of adventitious rooting assay, and pooled. Each biological replicate was formed by a pool of stem pieces collected from three different plants. Total RNA extraction and quantitative real-time PCR analyses were performed as previously described (Gutierrez *et al*., 2008) and are detailed in Methods S6. *PtUBIQUITINE* (Potri.001G418500), which had been previously validated for gene expression analysis in T89 stem cuttings with geNORM (Gutierrez *et al*., 2008) was used as the reference gene. Due to the high sequence similarity we failed to design paralogue-specific qPCR primers and instead designed primers that specifically amplify *PttARF6.1* and *PttARF6.2* paralogues together (*PttARF6.1/2*), *PttARF6.3* and *PttARF6.4* paralogues together (*PttARF6.3/4*). Similarly, primers were designed for *PttARF8.1* and *PttARF8.2* (*PttARF8.1/2*) and *PttARF17.1* and *PttARF17.2* (*PttARF17.1/2*) paralogue genes. Primers were designed to span the microRNA cleaving site for each gene to quantify the un-cleaved transcripts only (Table S1).

## Results

### Hybrid aspen and hybrid poplar show different patterns of adventitious root formation

To understand why some genotypes readily develop AR and others do not, we compared the rooting efficiency of cuttings from the poplar clone OP42 (*P. trichocarpa* × *P. maximowiczii*) and the hybrid aspen clone T89 (*P. tremula* × *P. tremuloides*) from juvenile plants kept *in vitro* (Fig. 1 and Fig. S1a,b,d) and from stem cuttings of three-month-old plants grown in the greenhouse (Fig. 2 and Fig. S1c,e). When cuttings were taken from juvenile *in vitro* plants, no significant difference between the two clones was observed (Fig. 1a). Nevertheless, in T89 *in vitro* cuttings, AR developed at the base of the cuttings in a crown-like arrangement (Fig. 1b-e), while in OP42, AR developed a few mm above the base of the cuttings and along the stem (Fig. 1 f-i,o,q). Cross- and longitudinal sections showed that in both cases the AR primordia initiated from the cambium region (Fig. 1j-q) as shown previously in cuttings of the *P. trichocarpa* clone 101-74 (Rigal *et al*., 2012). In contrast, when cuttings were taken from greenhouse-grown three-month-old plants (Fig. S1c) and kept in a hydroponic culture system as described elsewhere (Merret *et al*., 2010; Rigal *et al*., 2012) (Fig. S1e), T89 cuttings were unable to develop AR (Fig. 2a,b) while 100% of OP42 cuttings did root (Fig. 2a,c). For OP42 cuttings, the first bulges were visible on the stems as early as three days after cutting, and AR emerged after around five or six days (Fig. 2c) and fully developed and formed secondary roots were evident at 13 days after cutting (Fig. 2c). In both T89 and OP42 we observed the formation of lenticels; these correspond to cell proliferation regions in the cortex due to the high humidity in hydroponic conditions (Fig. 2b-e).

**Fig.1:**
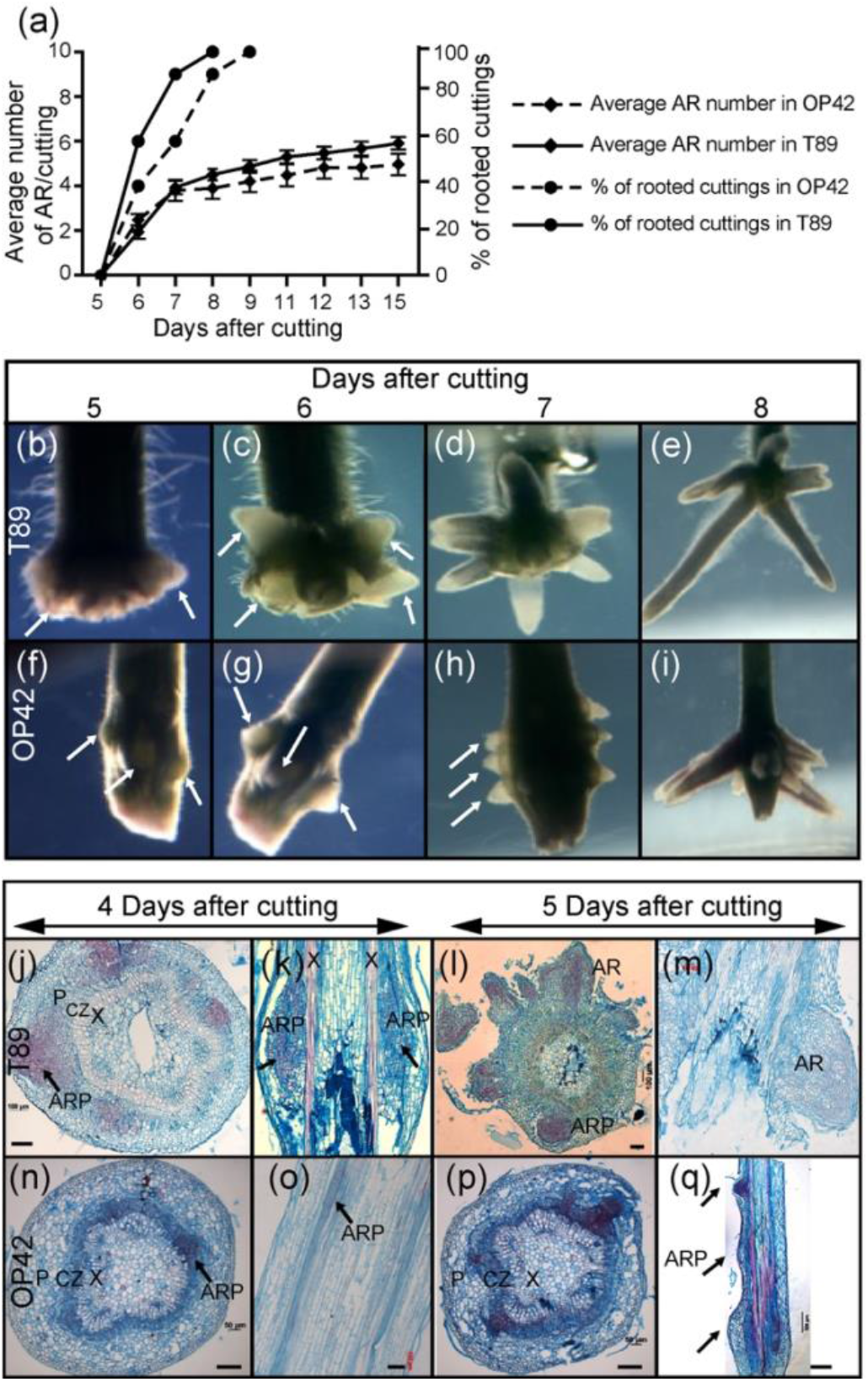
Pattern of adventitious rooting in hybrid aspen and hybrid poplar *in vitro*. (a) Average number of adventitious roots (AR) and percentage of rooted cuttings in T89 and OP42. Fifteen 3-cm-long cuttings, starting from the shoot apex, were taken from 4-week-old plantlets, amplified *in vitro*, and transferred onto half-strength MS medium as shown in Figs S1a,b,d). The emerged AR were scored starting on day 5 after transfer on fresh medium, until day 15. Data from three independent biological replicates, each of 15 stem cuttings, were pooled and averaged. Error bars indicate standard error. (b to e) In T89, AR developed all around the base of the cuttings in a crown-like formation as arrowed. (f to i) In OP42, AR developed few mm above the base of the cuttings and along the stem as arrowed. (j to q) Cross- (j, l, n, p) and longitudinal (k, m, o, q) sections show that in both cases the AR primordia develop from cells situated in the cambium/phloem region. CZ = cambial zone; P = Phloem; X = Xylem; APR = Adventitious root primordium; AR = Adventitious root.

**Fig.2:**
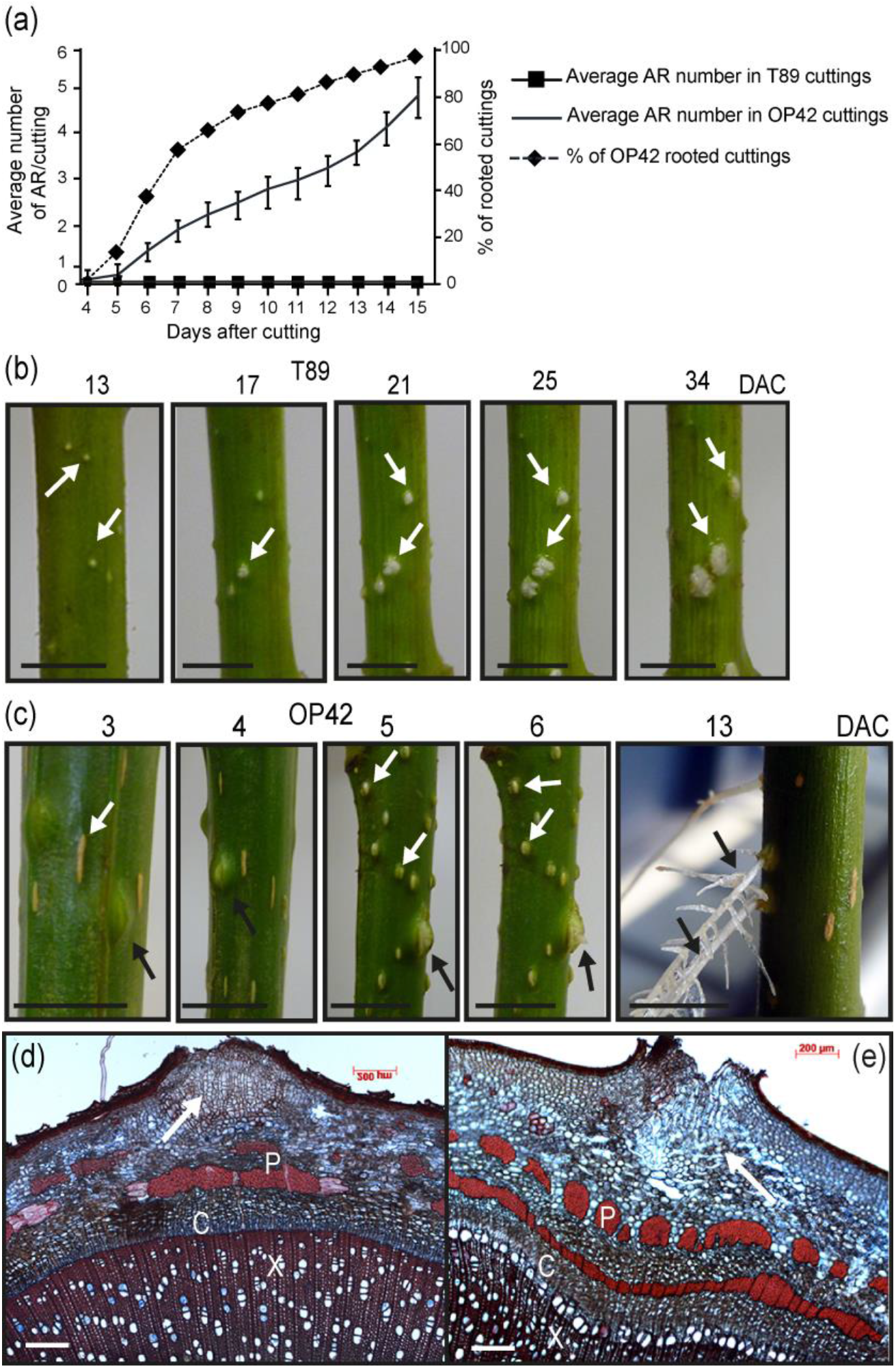
Adventitious root development in woody stem cuttings under hydroponic conditions. (a) Average number of adventitious roots (AR) and rooting percentage in T89 and OP42. About 20 cm lengths of stem from three-month-old greenhouse-grown hybrid aspen T89 and OP42 plants. The stem cuttings were kept in hydroponic conditions for five weeks and the number of AR was scored every day after cutting (DAC). Data from three biological replicates, each of at least 15 stem cuttings, were pooled and averaged. Error bars indicate standard error. (b) In T89 only lenticels were observed (white arrows). (c) In OP42, bulges of AR primordia were observed three DAC, and fully developed into AR 13 DAC (black arrows). Lenticels were also observed in OP42 cuttings (white arrows). (d, e) Cross-sections at the level of a lenticel (white arrows) in T89 (d) and OP42 (e). X = xylem; C = cambium; P = phloem.

### Transcriptomic profile and functional classification of Differentially Expressed Genes from cambium tissue between OP42 and T89 poplar genotypes

To explain this extreme difference in rooting performance, we performed a transcriptomic analysis of the cambium of OP42 and T89 cuttings from three-month-old plants grown in the greenhouse (Fig. S2a). We performed LCM (Fig. S2d-i) to dissect and collect homogenous and specific cambium tissue from the basal 5 mm of stem cuttings at time T0 (immediately after cutting) (Fig. S2b) and T1 (24 h after transfer in hydroponic conditions) (Fig. S2c). We mapped the RNA-seq reads to the *P. trichocarpa* reference genome (Dataset S1, sheet1) and classified 17,997 genes in the current annotation as being expressed significantly in all biological replicates in both genotypes at time T0 and T1 (Dataset S1, sheet 2). These 17,997 genes represent approximately 43% of the annotated genes in the *Populus* genome (poplar v3 assembly version; Tuskan *et al*., 2006). Interestingly, there were more DEGs in OP42 after 24 h in hydroponic conditions than in T89 (Fig. 3). In the case of T89, a total of 1198 (6.6% of the 17,997) genes were differentially expressed, 824 were up-regulated and 374 were down-regulated at T1 compared to T0 (Fig. 3a, Dataset S2, sheets 11 to 13). Gene Ontology (GO) enrichment analysis of DEGs showed a significant enrichment of GO terms related to biological processes, and molecular functions related to carbohydrate catabolism or redox mechanisms, regulation of transcription, response to abiotic stresses, cation binding, nucleic acid binding activity, or electron carrier activity (Dataset S3, sheets 4 and 5). In contrast, in OP42, a total of 5464 genes (30% of the 17,997 genes) were found differentially expressed, among which 3242 were up-regulated and 2222 down-regulated at time T1 compared to T0 (Fig. 3a,c. Dataset S2, sheets 8 to 10). Interestingly, among the 3242 DEGs, 2420 (74.6%) were exclusively up-regulated in OP42 at T1 (Fig. 3b), suggesting a specific remodulation of the transcriptome in OP42 during the 24 h timeframe that did not occur in T89. The GO enrichment analysis of these upregulated DEGs showed a significant enrichment of GO in cellular components, biological processes or molecular functions related to cell metabolism or cell biology such as transcription regulation, translation and post translation regulation (Dataset S3, sheet 4). Similarly, 66% of the 2222 DEGs that were down-regulated in OP42 at T1 compared to T0 were specifically differentially expressed in OP42 (Fig. 3c). In contrast to the up-regulated genes, the GO enrichment analysis showed a significant enrichment of GO in cellular components, biological processes or molecular functions related to abiotic stress responses (Dataset 3, sheet 5). When the two genotypes were compared to each other, 25% of the 17,997 genes were differentially expressed between OP42 and T89 at T0 (Fig. 3a, Dataset S2) among which, 2007 were up-regulated in T89 compared to OP42 (Fig. 3a) while 2533 were down-regulated (Fig. 3a, Dataset S2, sheets 2 to 4). This difference between the two genotypes was reduced to 14% 24 h after transfer into hydroponic conditions, with 1156 up-regulated and 1330 down-regulated in T89 compared to OP42 (Fig. 3a, Dataset S2, sheets 5 to 7). The genes that were differentially expressed between T89 and OP42 are mostly involved in cellular and chemical homeostasis, photosynthesis, dioxygenase activity and protein synthesis (Dataset S3, sheets 4 and 5).

**Fig.3:**
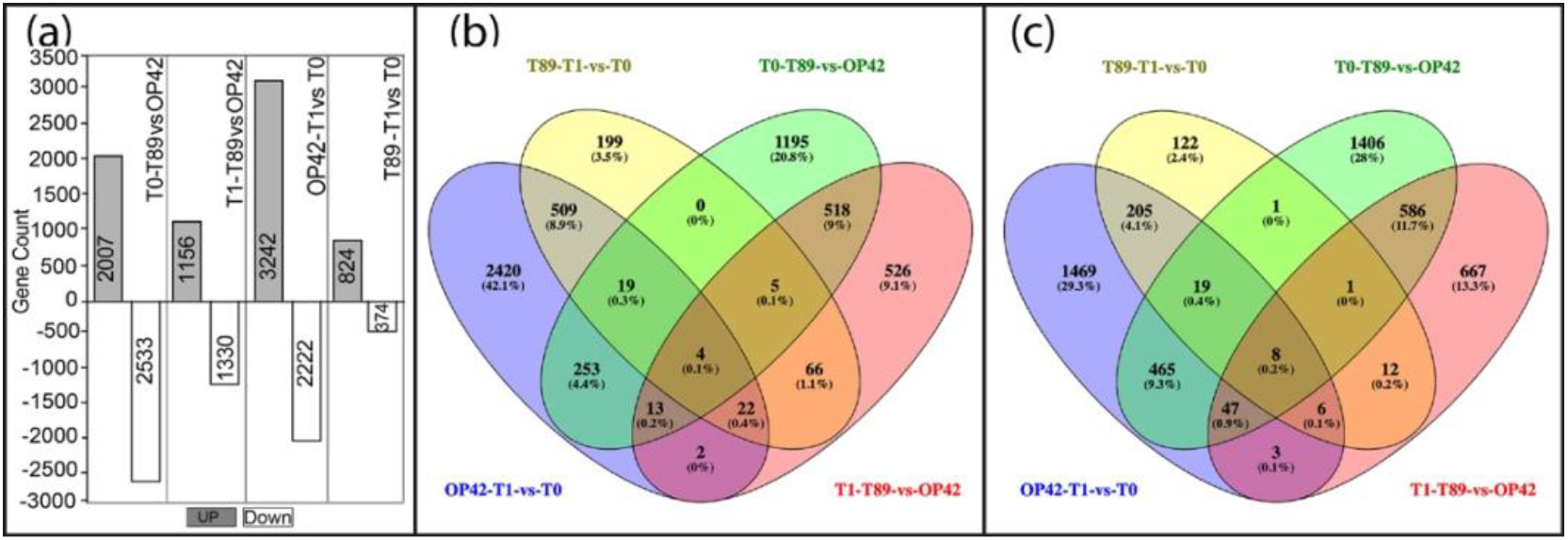
Number of differentially expressed genes (DEGs) between T89 and OP42. (a) Total number of differentially expressed genes up- and down-regulated in T89 and OP42. Venn diagram of DEGs between T89 and OP42 (b) Up-regulated (c) Down-regulated. Abbreviations signify as follows: T1-T89-vs-OP42; genes are up- or down-regulated in T89 compared to OP42 at time T1. T0-T89-vs-OP42; genes are up- or down-regulated in T89 compared to OP42 at time T0. T89-T1- vs-T0; genes are up- or down-regulated at time T1 compared to timeT0 in T89. OP42-T1-vs- T0; genes are up- or down-regulated at time T1 compared to time T0 in OP42.

### Genes related to cambium or vascular tissues behaved similarly in both genotypes

After checking the similarity of the biological replicates (Fig. S3a-b), we also confirmed the quality and the specificity of the Dataset. For this, we selected a list of 40 Arabidopsis genes described as being expressed in the cambium or vascular tissues, and checked the expression of their putative *Populus* orthologues in our Dataset (Fig. S3c and Dataset S3, sheet 1). All were found to be expressed (and most behaved similarly) in the two genotypes, showing a slight up-regulation or down-regulation in OP42 and T89 between T0 and T1 (Fig. S3c and Dataset S3, sheet 1). A few exceptions to this general pattern included Potri.003G111500 (*PtrPPNRT1.2*), Potri.004G223900 (similar to *AtCLAVATA1-related* gene) and Potri.014G025300 (similar to *AtWOX4b*) which were slightly down-regulated in T89 but up-regulated in OP42 24 h after cutting; additionally, a few genes were up-regulated in T89 compared to OP42 at T0 and T1. They comprise Potri.003G111500 (*PtrPPNRT1.2*), Potri.001G131800 (similar to Arabidopsis *BREVIS RADIX* gene) and Potri.002G024700 (ARF5), Potri.009G017700, which is similar to *AtLONESOME HIGHWAY*, a *bHLH* master transcriptional regulator of the initial process of vascular development.

### Genes encoding Reactive Oxygen Species scavenging proteins are mostly up-regulated in OP42 compared to T89

Reactive oxygen species (ROS) are signalling molecules involved in the response to biotic and abiotic stress as well as many aspects of plant development, including AR formation, as shown by recent studies (reviewed in Nag *et al*., 2013; Li *et al*., 2017; Velada *et al*., 2018). We therefore searched genes encoding ROS scavenging proteins among all DEGs in T89 and OP42. We identified 43 differentially expressed genes encoding ROS scavenging proteins, 33 of which belong to the GLUTATHIONE S-TRANSFERASE superfamily (GSTs) and ten to the PEROXIDASE superfamily (Dataset S3 sheet 3). Twenty of these genes were up-regulated at T1 compared to T0 in both genotypes, but on average the fold change was higher in OP42 than in T89 (Fig. S5; Dataset S3, sheet 3); nine genes were repressed 24 h after cutting in both genotypes. The most striking observation was that 32 out of 43 genes were significantly up-regulated in OP42 compared to T89 at T1, and 21 of those were also up-regulated in OP42 at T0 (Dataset S3, sheet 3); only six were up-regulated in T89 compared to OP42 at T0 and T1; four were up-regulated in T89 compared to OP42 at T0 but down-regulated in T89 compared to OP42 at T1; and five were up-regulated in OP42 compared to T89 at T0 - but by contrast - up-regulated in T89 at T1.

### The easy-to-root OP42 shows an increased transcriptional activity in the cambium compared to the difficult-to-root T89

The different stages of AR initiation (ARI) in *Populus* are associated with substantial remodelling of the transcriptome (Ramirez-Carvajal *et al*., 2009; Rigal *et al*., 2012). We therefore focused our analysis on the expression of transcription factors (TFs). From the 58 families of TF identified in *Populus*, 49 families were represented in the DEG list (Table 1; dataset S2; dataset S3, sheet 2) and most of the DEGs were observed in OP42 (Table 1). 24 h after cutting, 210 and 209 TF were up- or down-regulated respectively in OP42, while in T89 there were only 89 up-regulated and 43 down-regulated (Table 1). The most represented DEGs belong to the *ARF, bHLH*, bZIP, *C2H2*- and *C3H*-type zinc-finger family, *ERF, LBD, MYB, MYB-related, NAC* and *WRKY* families. Several genes belonging to those TF families have been shown to be involved in the control of adventitious rooting in *Populus* species (reviewed in Legue *et al*., 2014).

The *APETALA2/ETHYLENE RESPONSE FACTOR* (*AP2/ERF*) family was the most represented with 21 and 42 *ERF* genes up-regulated at T1 in T89 and OP42, respectively (Table 1 and Dataset S3, sheet 2). Twenty of the *ERFs* up-regulated in T89 were also up-regulated in OP42 24 h after cutting. Among the 22 specifically up-regulated in OP42, we found *PtrERF003* (Potri.018G085700; log_2_ FC = 7.7) (Dataset S3, sheet 2) which has been shown to be a positive regulator of AR development in *Populus* (Trupiano *et al*., 2013) and the *PtrERF39* (Potri.003G071700) a likely orthologue of the oxygen sensing *AtRAP2.12* (At1g53910) which has recently been shown to be involved in the primary root inhibition upon oxygen deficiency in Arabidopsis (Shukla *et al*., 2020). Several *WUSHEL-Like Homeobox* genes, have been shown to positively control AR development in *Populus* species (Li *et al*., 2018; Liu *et al*., 2014a; Liu *et al*., 2014b; Xu *et al*., 2015). More specifically, the *P. tomentosa PtoWOX5a* (Potri.008G065400) (Li *et al*., 2018), and the *Populus* × *euramericana PeWOX11/12ba* (Potri.013G066900) and *PeWOX11/12b* (Potri.019G040800) (Xu *et al*., 2015) have been found to be involved in AR development in poplar; nevertheless, they were not expressed in the cambium cells of OP42 or T89 (Dataset S1). By contrast, we found that two paralogues of *PtrWOX13, PtrWOX13a* (Potri.005G101800) and *PtrWOX13b* (Potri.005G252800) were up-regulated in OP42 24 h after cutting and transfer in hydroponic conditions (Dataset S3, sheet 2). *PtrWOX13* belongs to an ancient clade of *PtrWOX* genes (Liu *et al*., 2014b) and the Arabidopsis *AtWOX13* and *AtWOX14* are involved in the regulation of primary and lateral root development in Arabidopsis (Deveaux *et al*., 2008).

Recently (Wei *et al*., 2020) showed that the *P. ussuriensis PuHox52* gene, which belongs to the HD-Zip subfamily of TF is a positive regulator of adventitious rooting in *P. ussuriensis*. It acts by inducing nine regulatory hubs including the JA signalling pathway *PuMYC2* gene (MH644082; Potri.002G176900), a TF from the *bHLH* family, which has been demonstrated to be a positive regulator of AR development in *P. ussuriensis*. By contrast, in our dataset, we found that *P. trichocarpa PtrHox52* (Potri.014G103000) was down-regulated in the cambium of the easy-to-root genotype OP42 at T1 *i.e*., 24 h after cutting and transferred to hydroponic conditions (Dataset 3, sheet 2). *PtrHox52* was also up-regulated in the difficult-to-root genotype T89 compared to OP42 at T1 (Dataset S3, sheet 2). Accordingly, we observed that *PtrMYC2.5* (Potri.003G147300) was up-regulated in the cambium of T89 compared to OP42 at T1. There are six paralogues of *MYC2* in *Populus*. Three of them - *PtrMYC2.1* (Potri.003G092200), *PtrMYC2.2* (Potri.001G142200), *PtrMYC2.4* (Potri.001G083500) - were up-regulated in both T89 and OP42 at T1, but with a higher fold change in T89, while *PtrMYC2.5* (Potri.003G147300) was exclusively up-regulated in T89 at T1, which led to a significant increase in *PtrMYC2* expression in T89 compared to OP42 (Dataset S3, sheet 2). The potential up-regulation of JA signalling in T89 was corroborated by a higher fold change in the expression of several JA inducible *JA ZIM DOMAIN* (JAZ) genes 24 h after cutting in T89 compared to OP42. *PtrJAZ6* (Potri.003G068900), *PtrJAZ8* (Potri.011G083900) and *PtrJAZ10* (Potri.001G062500) were up-regulated in T89 compared to OP42 at T1 with a respective log2 FC of 4.25, 5.5 and 4.7 (Dataset S2, sheet 6). These results suggest a negative role of JA signalling on AR development as described in Arabidopsis (Gutierrez *et al*., 2012; Lakehal *et al*., 2020a) and contradict the positive role of JA on AR development as described for *P. ussuriensis* (Wei *et al*., 2020).

Several genes from the *AUXIN RESPONSE FACTOR* (*ARF*) have been shown to be involved in AR development in Arabidopsis and *Populus* (Gutierrez *et al*., 2009; Gutierrez *et al*., 2012; Lakehal *et al*., 2019; Cai *et al*., 2019; Liu *et al*., 2020). *AtARF6* and *AtARF8* are positive regulators of ARI while *AtARF17* negatively regulates adventitious rooting (Gutierrez *et al*., 2009). In *Populus, PeARF8* also positively regulates AR formation (Cai *et al*., 2019) but *PeARF17*, in contrast to the Arabidopsis gene, acts as a positive regulator of AR development in the hybrid poplar *P. davidiana* × *P. bolleana* (Liu *et al*., 2020). We identified 36 *PtrARF* genes encoding paralogues of 15 out of the 27 Arabidopsis *ARFs* orthologues. Although some of them were more significantly down-regulated in OP42 than in T89 24 h after cutting, they mostly behaved similarly in both genotypes (Fig. S6; Dataset S3, sheet 6). In particular, *PtrARF6.2* (Potri.002G055000) and *PtrARF6.3* (Potri.001G358500) were up-regulated while *PtrARF6.1* (Potri.005G207700) and *PtrARF6.4* (Potri.011G091900) were down-regulated in both T89 and OP42 at T1 compared to T0 (Fig. S6; Dataset S3, sheet 6). Similarly, both *PtrARF8.1* (Potri.004G078200) and *PtrARF8.2* (Potri.017G141000) were down-regulated at time T1 compared to T0 in both T89 and OP42. Interestingly, *PtrARF17.1* (Potri.002G089900) was significantly less expressed in the cambium of the difficult-to-root T89 than in OP42 at both time T0 and T1, which agrees with a potential positive role of *PtARF17.1* in AR development.

### *PttARF6* and *PttARF8* positively control adventitious rooting in hybrid aspen while *PttARF17* is a potential negative regulator

To assess the role of *PttARF6, PttARF8* and *PttARF17* in adventitious rooting in *Populus*, we produced transgenic plants that either over-expressed them or down-regulated their expression. Using the PopGenIE data base (http://popgenie.org) we identified the *Populus* genes most closely related to the Arabidopsis ones (Fig. S6a) and checked their expression pattern in the cambium and wood-forming region in the AspWood database (http://aspwood.popgenie.org/aspwood-v3.0/; Sundell *et al*., 2017). AspWood provides high resolution *in silico* transcript expression profiling of the genes expressed over the phloem, cambium, and other xylem development zones in aspen trees. We observed, *PtrARF6.1/2/3/4* and *PtrARF8.1/2* to be highly expressed in the phloem/cambium region while *PtrRF17.1/2* exhibited very low expression in the same region (Fig. S4B-D).

For the over-expressing lines *PttARF6.4* and *PttARF8.1*, coding sequences were cloned under the control of the 35S promoter of the Cauliflower Mosaic Virus (CaMV) or the promotor of the cambium specific gene *PtrHB3a* (Schrader *et al*., 2004). For down-regulated lines RNAi constructs were made to target *PttARF6.3* and *4*, *PttARF8.1* and *2*, and *PttARF17.1* and *2* paralogues. We had previously shown that in Arabidopis hypocotyl, *AtARF6*, *AtARF8* and *AtARF17* regulate each other’s expression at the transcriptional and post-transcriptional level and that the balance between positive and negative regulators determined the average number of AR (Gutierrez *et al*., 2009). As in Arabidopsis, the *Populus ARFs* are regulated by microRNAs (Cai *et al*., 2019; Liu *et al*., 2020). We therefore checked the relative transcript amount of the un-cleaved transcript of the three *ARF* types in each transgenic line. A multiple sequence alignment analysis revealed that the coding sequences (CDS) of *PttARF6.1* and *PttARF*6.2 paralogues were highly similar, and we were unable to differentiate their expression level by q-PCR. A similar situation occurred with *PttARF6.3* and *PttARF6.4, PttARF8.1* and *Ptt*A*RF*8.2, *PttARF17.1* and *PttARF17.2*., We therefore designed primers to span the microRNA cleaving site and measured the cumulative expression level of the two paralogues (designated *PttARF6_1*+*2*; *PttARF2_3*+*4*; and *PttARF17_1*+*2*) (Fig. 4 and Fig. S7a, b).

**Fig. 4:**
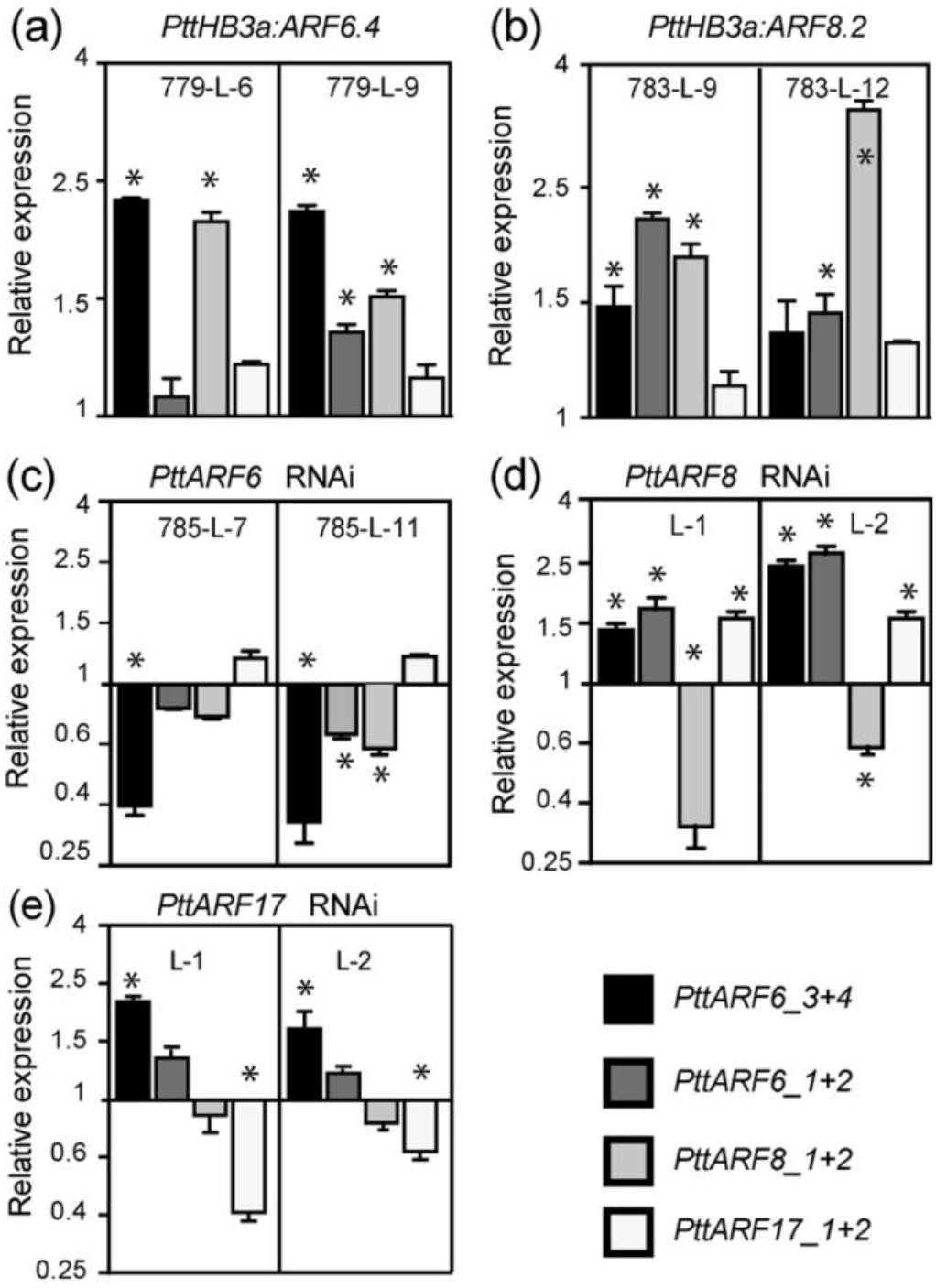
Relative un-cleaved transcript amount of *PtARF6.1/2, PtARF6.3/4, PtARF8.1/2, PtARF17.1/2* in transgenic lines overexpressing or downregulated for *PtARF6, PtARF8* or *PtARF17*. The *PtARF6.1/2, PtARF6.3/4, PtARF8.1/2, PtARF17.1/2* un-cleaved transcript abundance was quantified in stem cutting fragments of independent over-expressing (a, b) or down-regulated (c-e) lines. Gene expression values are relative to the reference genes and calibrated toward the expression in the control line T89, for which the value is set to 1. Error bars indicate SE obtained from three independent biological replicates. A one-way analysis of variance combined with the Dunnett’s comparison post-test indicated that the values marked with an asterisk differed significantly from T89 values (P < 0.05; n = 3).

We confirmed the over-expression of *PttARF6_3*+*4* and *PttARF8_1*+*2* in the over-expressing lines (Fig. 4a,b and Fig. S7a,b), and the down-regulation of *PttARF6_3*+*4*, *PttARF8_1*+*2* and *PttARF17_1*+*2* in the RNAi lines (Fig. 4c-e). Interestingly, we observed that, as in Arabidopsis, when the expression of one of the three ARFs was modified, the expression of the others was also affected, establishing a different ratio between potential positive and negative regulators (Fig. 4 and Fig. S7).

We performed rooting assays to check the aptitude of the different transgenic lines in producing AR. When either *PttARF6.4* or *PttARF8.2* was over-expressed in the cambium under the control of the *PttHB3* promoter, the transgenic lines produced more AR than the control T89 (Fig. 5a,b). Similar results were obtained with *PttARF6.4* under the 35S promotor (Fig. S7c) but not with *p35SPttARF8.2* (Fig. S7d). The positive effect of *PttARF6* and *PttARF8* was confirmed in the RNAi lines, which produced fewer AR than the control line T89 (Fig. 5c,d). The role of *PttARF17* was unclear, although it has been described as a positive regulator in the hybrid poplar *P. davidiana* × *P. bolleana* (Liu *et al*., 2020). However, our results show that when *PttARF17_1*+*2* is down-regulated the transgenic lines produce more AR (Fig. 5E) suggesting that *PttARF17.1* or *PttARF17* could be negative regulators. Nevertheless, because *PttARF6_3+4* were up-regulated in the *PttARF17* RNAi lines (Fig. 4e), it is difficult to conclude whether the increased AR average number was solely due to the down-regulation of *PttARF17*, to the over-expression of *PttARF6_3*+*4*, or to a combination of both.

**Fig. 5:**
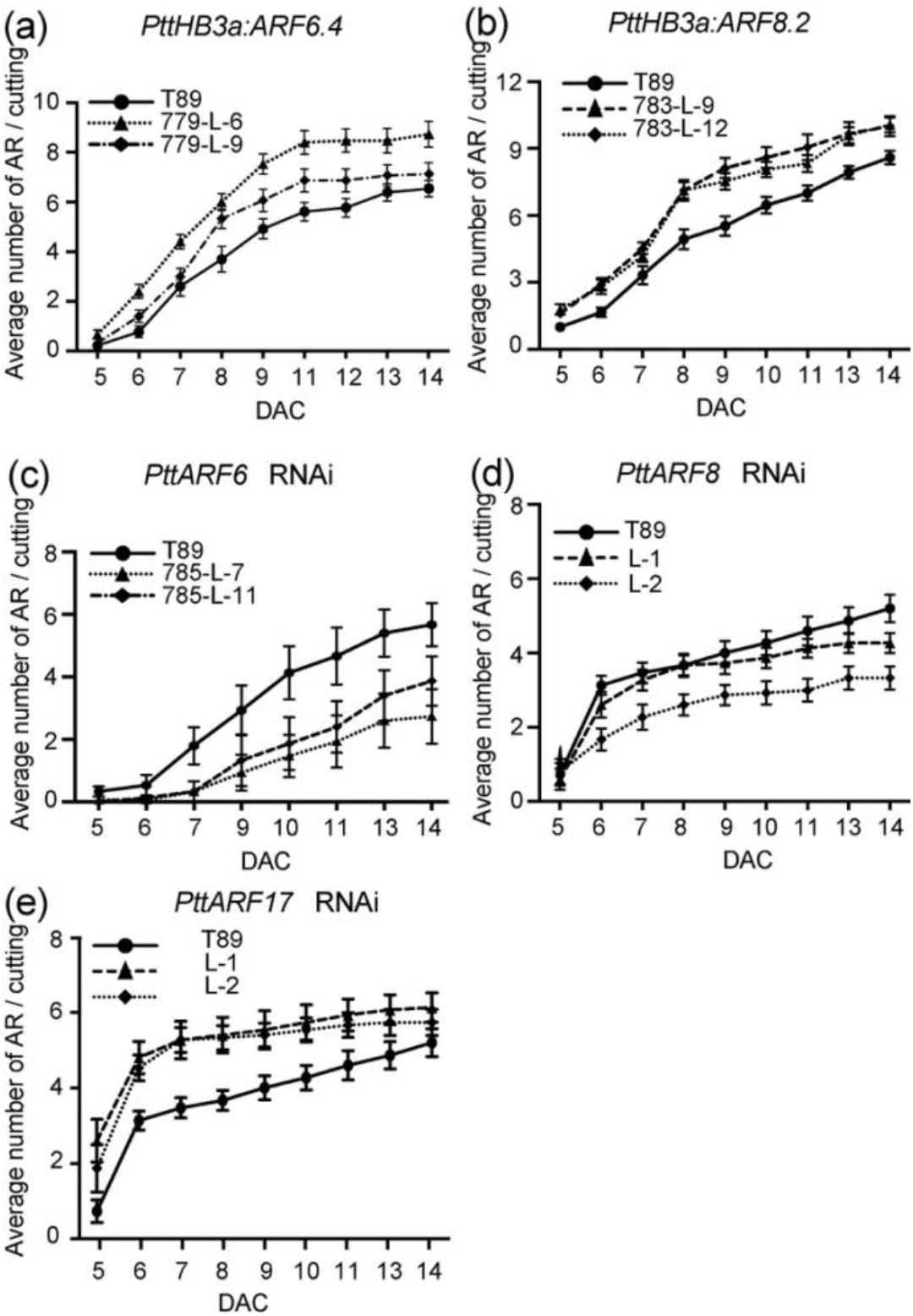
*PtARF6* and *PtARF8* positively control adventitious root (AR) development while *PtARF17* is a negative regulator. (a, b) Average number of AR on cuttings of transgenic plants expressing *PtARF6.4* (a) and *PtARF8.2* (b) under the cambium specific promoter *pPtHB3*. Rooting assay was performed as described in Material and Methods. Two independent transgenic lines were compared to the control T89. AR number was scored every day starting day 5 after cutting until 14 days after cut (DAC). For each line 15 cuttings were analysed. (c-e) Average number of AR on cuttings of transgenic plants expressing the *p35S:PtARF6.2-RNAi* (c), *p35S:PtARF8.4-RNAi* (d) or *p35S:PtARF17.2-RNAi* (e) constructs. Two independent transgenic lines were compared to the T89 control. AR number was scored every day starting day 5 after cutting until 14 DAC. For each line 15 cuttings were analysed. Data are means ± SE, n = 15, corresponding to two independent lines per construct. A two-way ANOVA with a Tukey’s multiple comparisons test indicated that the difference between the transgenic lines and the control were significant, except for *PtHB3a:ARF6.4* line 779-L-9 for which the difference was significant only from day 8 to 12, and *PtARF8-RNAi* L-1 for which no significant difference was observed.

### *PtMYC2.1* is a negative regulator of adventitious root development in hybrid aspen

In Arabidopsis, the *AtARF6, AtARF8* and *AtARF17* have been shown to act upstream of *AtMYC2*, which is a negative regulator of AR development (Gutierrez *et al*., 2012; Lakehal *et al*., 2020a). In our present study, five out of the six *PtrMYC2* paralogues are shown to be among the DEGs (Fig. 6a, Dataset S3, sheet 2). They mostly behaved the same way in both T89 and OP42, but the fold change induction was higher for four of them at T1 in the difficult-to-root genotype T89, and *PtMYC2.5* was significantly up-regulated in T89 compared to OP42 24 h after cutting (Fig. 6a, Dataset S3, sheet 2). These results suggest that *PtrMYC2* could be a negative regulator of adventitious rooting in hybrid aspen. To confirm this hypothesis, we produced transgenic hybrid aspen trees over-expressing *PttMYC2.1* under the 35S promoter. The over-expression was confirmed in two independent transgenic lines by qPCR (Fig. 6b) and the rooting assays confirmed that over-expressing *PttMYC2.1* repressed AR development (Fig. 6c). The up-regulation of the JA signalling pathway in T89 cambium compared to OP42 could contribute to the recalcitrance of stem cuttings from greenhouse-grown plants to produce AR. This led us to compare the behaviour of OP42 and T89 in response to exogenous JA. Rooting assays were performed with *in vitro* propagated T89 and OP42 plants in the absence or presence of increasing concentrations of JA (Fig. 6 c,d). We observed that even though the two genotypes root similarly under *in vitro* conditions, they showed a different response to exogenous JA. The difficult-to-root T89 was more sensitive to exogenously applied JA compared to OP42 (Fig. 6 c,d)

**Fig. 6:**
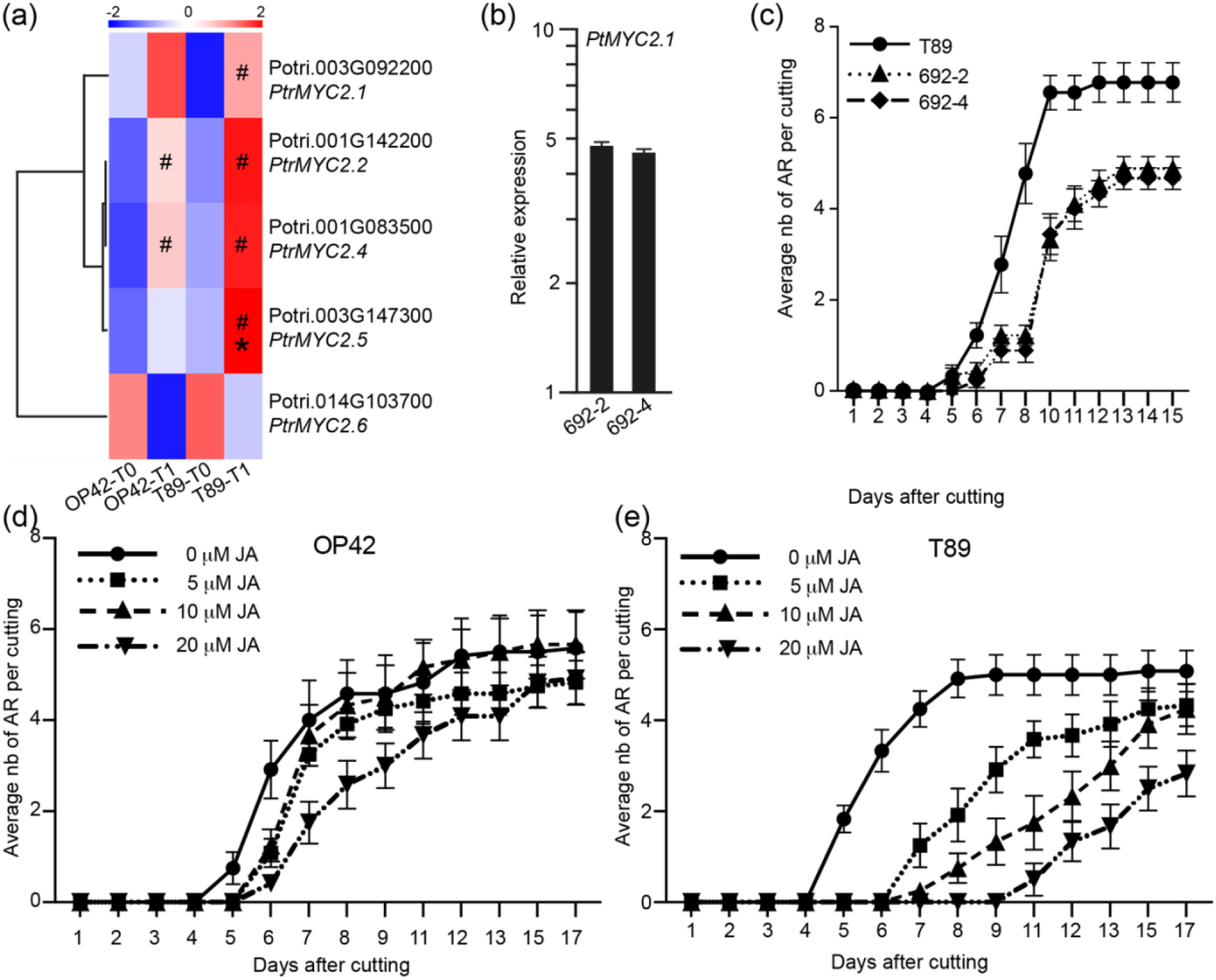
Jasmonate is a negative regulator of AR development in hybrid aspen cuttings. (a) The expression of five out of six *PtMYC2* paralogues found in the transcriptomic data set presented as a heat map clustering in T89 and OP42 at time T0 and T1. Colours indicate low expressed genes (blue) or highly expressed genes (red) (b) *PtMYC2.1* transcript abundance was quantified in stem cutting fragments of two independent transgenic T89 lines over-expressing *PtMYC2.1* under the 35S promotor (lines 692-2 and 692-3). Gene expression values are relative to the reference gene and calibrated toward the expression in the control line T89, for which the value is set to 1. Error bars indicate SE obtained from three independent biological replicates. (c) Average number of AR in stem cuttings of over-expressing *PtMYC2.1* transgenic T89 compared to the wild type T89. For each line 15 cuttings were analysed. Data are means ± SE, n = 15. (d-e) Average number of AR in stem cuttings of (c) OP42 and (d) T89 in the absence or presence of 5μM, 10μM and 20μM methyl jasmonate. For each line and each condition 15 cuttings were analysed. Data are means ± SE, n = 15. Three independent biological replicates were used. A two-way ANOVA with a Tukey’s multiple comparisons test indicated that: In the case of OP42 a significant difference between non treated plants and treated plants was observed at day 6 for all JA concentrations (P < 0.05 for 5 and 10 μM JA, P < 0.0001 for 20 μM JA) and then at day 7 and 8 only in presence of 20 mM JA (P < 0.01). For T89 a very significant effect of JA was observed for all concentrations from day 5 until day 15 (P < 0.0001 for 10 and 20 μM, P < 0.05 from day 5 until day 12)

## Discussion

*Populus* species are among the most economically utilised trees. Their ability to be propagated vegetatively means that novel genotypes can be rapidly multiplied. Nevertheless, tree cloning is often limited by the difficulty of developing AR from stem cuttings. Adventitious rooting is a complex multifactorial process. Many QTL have been detected for adventitious rooting-related traits (Ribeiro *et al*., 2016; Sun *et al*., 2019; Zhang *et al*., 2009) highlighting the genetic complexity of this trait. With the emergence of Arabidopsis as a genetic model, many genes and signalling pathways involved in the control of AR development have been identified (Gutierrez *et al*., 2009; Gutierrez *et al*., 2012; Hu and Xu, 2016; Lakehal *et al*., 2019; Lakehal *et al*., 2020a; Lakehal *et al*., 2020b; Liu *et al*., 2014b; Sorin *et al*., 2005), and lately, several groups have focused on AR development in *Populus* and identified genes and gene networks involved in this process (Cai *et al*., 2019; Legue *et al*., 2014; Li *et a*l., 2018; Ramirez-Carvajal *et al*., 2009; Trupiano *et al*., 2013; Wei *et al*., 2020; Xu *et al*., 2021; Xu *et al*., 2015; Yordanov *et al*., 2017; Yue *et al*., 2020; Zhang *et al*., 2020). Nevertheless, despite all this research, most has so far focused on successive AR development stages in a given genotype; there have been no comparisons between easy-to-root and difficult-to-root genotypes.

To understand the underlying causes of poor-rooting and good-rooting in different genotypes we compared the hybrid poplar clone OP42, which is easily propagated from dormant stem cuttings, and the hybrid aspen clone T89, which is unable to develop AR under the same conditions.

Previous research has revealed that, predictably, AR form from specific founder cells in poplar stem cuttings, but that the process is highly dependent upon induction treatment and age of the cutting (Rigal *et al*., 2012). Cambium cells have also been shown to be competent initiators of AR in *Eucalyptus* or *Populus* (Chao *et al*., 2019; Chiatante *et al*., 2010). Transcriptomic profiling of vascular tissues including the cambium region in *Populus* have been reported in several studies (Schrader, et al., 2004; de Almeida *et al*., 2015; Kim *et al*., 2019), but little attention has been given to gene expression in *Populus* cambial cells during AR development. Rigal *et al*. (2012) showed that changes in the transcriptome occur in the cambium during the early stages of AR development in *Populus*. In our present study we performed a global comparative transcriptomic analysis of the cambium of cuttings taken from OP42 and T89 clones.

Interestingly, the juvenile plants from the two clones rooted similarly when grown *in vitro*. In both cases the AR originate from the cambium region. But the hybrid aspen T89, unlike the hybrid poplar OP42, was unable to develop roots from 3-month-old plants grown in the greenhouse. Aging is a well-known limiting factor for AR development (reviewed in Aumond Jr. *et al*., 2017; Bellini *et al*., 2014; Diaz-Sala *et al*., 2002). What cellular and biochemical modifications occur during maturation and phase changes and how these events reconfigure molecular pathways that lead to the inhibition of ARI in mature tissues is still not well understood. A comparison of DNA methylation in samples from juvenile and mature chestnut cuttings has shown that aging is related to a progressive increase of methylated 5-deoxycytidines (Baurens *et al*., 2004; Hasbun *et al*., 2007; Monteuuis *et al*., 2008). In contrast, progressive reduction in DNA methylation by grafting of adult shoot scions of coast redwood *(Sequoia sempervirens)* onto juvenile rootstock resulted in the progressive restoration of juvenile traits and rooting competence (Huang *et al*., 2012). The connection between phase changes and epigenetic gene regulation has been confirmed by the fact that several Arabidopsis mutants affected in phase change were also altered in the genesis of small RNAs (19–24-nucleotide RNAs), including both microRNAs and short interfering RNAs (Willmann and Poethig, 2005), and microRNA *miR156*, which, as one of the most evolutionally conserved miRNAs in plants, is one of the regulators of the ‘age pathway’ (reviewed in Wang, 2014).

Congruent with a potential effect of age-related mechanisms on gene expression, we observed that there were many more DEGs in OP42 than in T89, 24 hours after cutting and transfer to rooting conditions. There were many more transcription factors differentially expressed in OP42 suggesting a more extensive transcriptome reprogramming activity in the cambium during the early stages of ARI.

Interestingly, among the differentially expressed transcription factors we found that the *P. trichocarpa PtHox52* gene (Potri.014G103000) was down-regulated in the cambium of the easy-to-root genotype OP42 and up-regulated in the difficult-to-root genotype T89 compared to OP42 at T1. This is surprising since the *P. ussuriensis PuHox52* gene, has been described as a positive regulator of adventitious rooting in *P. ussuriensis* (Wei *et al*., 2020). It was shown to induce nine regulatory hubs including the JA signalling pathway driven by the *PuMYC2* gene (MH644082; Potri.002G176900), which was confirmed to be a positive regulator of AR development in *P. ussuriensis*. By contrast, JA signalling appears to be up-regulated in the cambium of the difficult-to-root T89 genotype compared to OP42, and we confirmed that *PtMYC2.1* negatively controls AR development in the hybrid aspen T89 as we had previously shown in Arabidopsis (Gutierrez *et al*., 2012; Lakehal *et al*., 2020a). These are intriguing results, but the role of JA in the control of AR is still not totally clear and seems to be context and species dependent (Lakehal *et al*., 2020b). It will be interesting in the future to study whether *Populus MYC2* paralogues have acquired different functions depending on the species, the growth and vegetative propagation conditions. Although T89 and OP42 clones rooted similarly *in vitro*, T89 was more sensitive to exogenously applied JA. This result suggests that the higher up-regulation of the JA pathway in the cambium of T89 24 h after cutting could contribute to repress ARI.

Interestingly, the orthologues of the three Arabidopsis *ARF* genes that were shown to be either positive (*AtARF6*, *AtARF8*) or negative (*AtARF17*) regulators of ARI in Arabidopsis (Gutierrez *et al*., 2009; Gutierrez *et al*., 2012; Lakehal *et al*., 2019) behaved similarly in both T89 and OP42. An exception is *PttARF17.1*, which was significantly less expressed in the cambium of the difficult-to-root T89 as compared to OP42 at both time points T0 and T1. This result agrees with a potential positive role of *PttARF17.1* in ARI as described for *PeARF17* in the hybrid poplar *P. davidiana* × *P. bolleana* (Liu *et al*., 2020). Nevertheless, a down-regulation of *PttARF17.1* and *PttARF17.2* expression in T89 induced ARI suggesting a negative role for *PttARF17*. As in Arabidopsis (Gutierrez *et al*., 2009) when the expression of one of the three *PttARFs* was perturbed, the expression of the others was modified. In our current case of the down-regulation of *PttARF17*, *PttARF6* paralogues were up-regulated, which probably contributed to increase ARI. As for *MYC2* genes, it is possible that different paralogues of *ARF17* have different functions depending on the species or the context.

There were many transcription factors that were either up- or down-regulated in OP42 at T1 compared to T0 but not in T89, and their further characterisation may certainly further advance our understanding of the mechanisms differentiating difficult-to-root from easy-to-root genotypes.

Another interesting difference we observed between T89 and OP42 concerns the expression of genes encoding ROS scavenging proteins. We identified 43 of these genes among the DEGs, 33 of which belong to the GST superfamily and 10 to the PEROXIDASE superfamily. The most striking observation was that 32 were significantly up-regulated in OP42 compared to T89 at T1, and 21 of those were also up-regulated in OP42 at T0. Recent studies have shown that peroxidase activity positively regulates AR formation in different species (reviewed in Nag *et al*., 2013; Li *et al*., 2017; Velada *et al*., 2018). It is therefore possible that the up-regulation of most of these genes in the cambium of OP42 compared to T89 partially explains the difference in rooting competence.

## Supporting information

Supplemental dat set 1

Supplemental data set 2

Supplemental data set 3

## Acknowledgments

The authors sincerely thank Dr Marta Derba Maceluch from the UPSC Microscopy platform, and Dr Nicolas Delhomme and the UPSC bioinformatic platform for their support with the data analysis; the personnel from the UPSC transgenic facility for the production of the transgenic plants; and Dr Didier Le Thiec from INRAE of Nancy for the use of cryostat and his help and useful suggestions.

## Author contributions

AR, IP, SA, VL and CB conceived and designed the experiments. AR, IP, SA, RS, AK and FB performed the experiments. AR and CB wrote the manuscript. SA, IP reviewed and edited the manuscript. All authors read, commented and approved the final article for publication. RB, VL and CB supervised the work. SA, FM, AK, RB and CB acquired funding.

## Data availability

The RNA-seq data have been deposited at the European Nucleotide Archive (http://www.ebi.ac.uk/ena/) and will be available using the accession number PRJEB21558 (OP42 and T89 data under PRJEB21549 and PRJEB21557, respectively).

**The following Supporting Information is available for this article:**

**Methods S1** Plant growth conditions and rooting assays

**Methods S2** Histological analysis of stem cuttings *in vitro*

**Methods S3** Tissue preparation before laser capture microdissection

**Methods S4** Pre-processing of RNA-Seq data

**Methods S5** Generation of plasmid constructs and transformation of hybrid aspen

**Methods S6** Quantitative Real-Time PCR analysis

**Fig. S1** Conditions for adventitious rooting assays from *in vitro* plants and greenhouse-grown plants

**Fig. S2:** Workflow for laser capture microdissection (LCMS) of cambium tissues from stem **cuttings**

**Fig. S3**: Quality assessment of the RNAseq data in the different biological replicates

**Fig. S4**: Populus Arabidopsis orthologues of *ARF6, ARF8* and *ARF17* and their expression pattern in wood-forming tissues

**Fig.S5:** Heat map showing the average expression of genes encoding ROS scavenging proteins in the cambium of T89 and OP42 genotypes

**Fig.S6:** Heat map showing the average expression of *PtrARF* genes in the cambium of T89 and OP42 genotypes

**Fig. S7: Over-expression** of *PtAF6.4* and PtARF8.2 under the 35S promoter

**Table S1** Primer list used in the present study.

**Dataset S1** (excel document):

**Sheet 1** Library-size-normalised variance-stabilised data set

**Sheet 2** Expression values for the 17,997 expressed genes

**Dataset S2:** Differentially expressed genes

**Sheet 1** The differentially expressed genes (DEG) up- and down-regulated and their annotation.

**Sheet** 2 Total number of DEG in OP42 when compared at time T1 and T0.

**Sheet 3** The DEG up-regulated in OP42 at time T1 compared to time T0.

**Sheet 4** The DEG down-regulated in OP42 at time T1 compared to time T0.

**Sheet 5** Total number of DEG in T89 when compared at time T1 and T0.

**Sheet 6** The DEG up-regulated in T89 at time T1 compared to time T0.

**Sheet 7** The DEG down-regulated in T89 at time T1 compared to time T0.

**Sheet 8** The total number of DEG in OP42 and T89 at time T0.

**Sheet 9** The number of DEG up-regulated in T89 compared to OP42 at timeT0.

**Sheet 10** The number of DEG down-regulated in T89 compared to OP42 at timeT0.

**Sheet 11** Total number of DEG between T89 and OP48 at time T1.

**Sheet 12** The number DEG up-regulated in T89 compared to OP42 at time T1.

**Sheet 13** The number of DEG down-regulated in T89 compared to OP42 at time T1.

**Dataset S3**

**Sheet 1** Vascular tissue expressed genes set.

**Sheet 2** Differentially expressed Transcription Factors.

**Sheet 3** Differentially expressed ROS scavenging proteins

**Sheet 4** Gene Ontology of up-regulated DEGs.

**Sheet 5** Gene Ontology of down-regulated DEGs.

**Sheet 6** Mean of expression values used for the heat maps

## References

Abarca, D., and Díaz-Sala, C. 2009a. Adventitious root formation in conifers. In: Adventitious Root Formation of Forest Trees and Horticultural Plants – from Genes to Applications. Eds: Niemi, K., and Scagel, C., eds. Kerala, India: Research Signpost Publishers. 227.

Abarca, D., and Díaz-Sala, C. 2009b. Reprogramming adult cells during organ regeneration in forest species. Plant Signal Behav 4:793–795.

Aumond Jr., M.L., de Araujo Jr., A.T., de Oliveira Junkes, C.F., De Almeda, M.R., Matsuura, H.N., de Costa, F., and Fett-Neto, A.G. 2017. Events Associated with Early Age-Related Decline in Adventitious Rooting Competence of Eucalyptus globulus Labill. Frontiers in Plant Science 8:1734.

Bannoud, F. and Bellini, C. (2021) Adventitious Rooting in Populus Species: Update and Perspectives. Frontiers in Plant Science 12:668837

Baurens, F.C., Nicolleau, J., Legavre, T., Verdeil, J.L., and Monteuuis, O. 2004. Genomic DNA methylation of juvenile and mature Acacia mangium micropropagated in vitro with reference to leaf morphology as a phase change marker. Tree Physiol 24:401–407.

Bellini, C., Pacurar, D.I., and Perrone, I. 2014. Adventitious roots and lateral roots: similarities and differences. Annu Rev Plant Biol 65:639–666.

Bozzano, M., Jalonen, R., Thomas, E., Boshier, D., Gallo, L., Cavers, S., Bordács, S., Smith, P., and Loo, J. 2014. Genetic considerations in ecosystem restoration using native tree species. State of the World’s Forest Genetic Resources – Thematic Study. Rome: FAO and Bioversity International.

Brunoni, F., Ljung, K., and Bellini, C. 2019. Control of root meristem establishment in conifers. Physiol. Plant. 165: 81–89.

Cai, H., Yang, C., Liu, S., Qi, H., Wu, L., Xu, L.A., and Xu, M. 2019. MiRNA-target pairs regulate adventitious rooting in *Populus*: a functional role for miR167a and its target Auxin response factor 8. Tree Physiol 39:1922–1936.

Carle, J., Ball, J.B., and del Lungo, A. 2008. The global thematic study of planted forests. In: Planted forests: uses, impacts and sustainability. --Evans, J., ed. Wallingford (UK) and Rome (Italy): CABI and FAO. 33–46.

Chao, Q., Gao, Z.F., Zhang, D., Zhao, B.G., Dong, F.Q., Fu, C.X., Liu, L.J., and Wang, B.C. 2019. The developmental dynamics of the *Populus* stem transcriptome. Plant Biotechnol J 17:206–219.

Chiatante, D., Beltotto, M., Onelli, E., Di Iorio, A., Montagnoli, A., and Scippa, S.G. 2010. New branch roots produced by vascular cambium derivatives in woody parental roots of *Populus nigra*. Plant Biosystems 144:420–433.

de Almeida, M.R., de Bastiani, D., Gaeta, M.L., de Araujo Mariath, J.E., de Costa, F., Retallick, J., Nolan, L., Tai, H.H., Stromvik, M.V., and Fett-Neto, A.G. 2015. Comparative transcriptional analysis provides new insights into the molecular basis of adventitious rooting recalcitrance in Eucalyptus. Plant Sci 239:155–165.

Deveaux, Y., Toffano-Nioche, C., Claisse, G., Thareau, V., Morin, H., Laufs, P., Moreau, H., Kreis, M., and Lecharny, A. 2008. Genes of the most conserved WOX clade in plants affect root and flower development in Arabidopsis. BMC Evol Biol 8:291.

Diaz-Sala, C., Garrido, G., and Sabater, B. 2002. Age-related loss of rooting capability in Arabidopsis thaliana and its reversal by peptides containing the Arg-Gly-Asp (RGD) motif. Physiol Plant 114:601–607.

Dickmann, D.I. 2006. Silviculture and biology of short-rotation woody crops in temperate regions: then and now. Biomass Bioenergy 30:696–705.

Dobin, A., Davis, C.A., Schlesinger, F., Drenkow, J., Zaleski, C., Jha, S., Batut, P., Chaisson, M., and Gingeras, T.R. 2013. STAR: ultrafast universal RNA-seq aligner. Bioinformatics 29:15–21.

Geiss, G., Gutierrez, L., and Bellini, C. 2009. Adventitious root formation: new insights and perspective. In: Root Development - Annual Plant Reviews --Beeckman, T., ed. London: Blackwell Publishing-CRC Press. 127–156.

Gentleman, R.C., Carey, V.J., Bates, D.M., Bolstad, B., Dettling, M., Dudoit, S., Ellis, B., Gautier, L., Ge, Y., Gentry, J., et al. 2004. Bioconductor: open software development for computational biology and bioinformatics. Genome Biol 5:R80.

Gou, J., Strauss, S.H., Tsai, C.J., Fang, K., Chen, Y., Jiang, X., and Busov, V.B. 2010. Gibberellins regulate lateral root formation in *Populus* through interactions with auxin and other hormones. Plant Cell 22:623–639.

Gutierrez, L., Bussell, J.D., Pacurar, D.I., Schwambach, J., Pacurar, M., and Bellini, C. 2009. Phenotypic plasticity of adventitious rooting in Arabidopsis is controlled by complex regulation of AUXIN RESPONSE FACTOR transcripts and microRNA abundance. Plant Cell 21:3119–3132.

Gutierrez, L., Mauriat, M., Guenin, S., Pelloux, J., Lefebvre, J.F., Louvet, R., Rusterucci, C., Moritz, T., Guerineau, F., Bellini, C., and Van Wuytswinkel, O. 2008. The lack of a systematic validation of reference genes: a serious pitfall undervalued in reverse transcription-polymerase chain reaction (RT-PCR) analysis in plants. Plant Biotechnol J 6:609–618.

Gutierrez, L., Mongelard, G., Flokova, K., Pacurar, D.I., Novak, O., Staswick, P., Kowalczyk, M., Pacurar, M., Demailly, H., Geiss, G., et al. 2012. Auxin controls Arabidopsis adventitious root initiation by regulating jasmonic acid homeostasis. Plant Cell 24:2515–2527.

Hamann, T., Smets, E., and Lens, F. 2011. A comparison of paraffin and resin-based techniques used in bark anatomy. Taxon 60:841–851.

Hasbun, R., Valledor, L., Santamaria, E., Canal, M.J., and Rodriguez, R. 2007. Dynamics of DNA methylation in chestnut trees development. In: Proc. 27th IHCS-S1 Plant Gen. Ressources--Hummer, K.E., ed.: Acta Hort. 760. 563–566.

Hu, X., and Xu, L. 2016. Transcription Factors WOX11/12 Directly Activate WOX5/7 to Promote Root Primordia Initiation and Organogenesis. Plant Physiol 172:2363–2373.

Huang, L.C., Hsiao, L.J., Pu, S.Y., Kuo, C.I., Huang, B.L., Tseng, T.C., Huang, H.J., and Chen, Y.T. 2012. DNA methylation and genome rearrangement characteristics of phase change in cultured shoots of Sequoia sempervirens. Physiologia plantarum 145:360–368.

Karlberg, A., Bako, L., and Bhalerao, R.P. 2011. Short day-mediated cessation of growth requires the down regulation of AINTEGUMENTALIKE1 transcription factor in hybrid aspen. PLoS Genet 7:e1002361.

Kim, M.H., Cho, J.S., Jeon, H.W., Sangsawang, K., Shim, D., Choi, Y.I., Park, E.J., Lee, H., and Ko, J.H. 2019. Wood Transcriptome Profiling Identifies Critical Pathway Genes of Secondary Wall Biosynthesis and Novel Regulators for Vascular Cambium Development in *Populus*. Genes (Basel) 10.

Kirilenko, A.P., and Sedjo, R.A. 2007. Climate change impacts on forestry. Proc Natl Acad Sci U S A 104:19697–19702.

Kucukoglu, M., Nilsson, J., Zheng, B., Chaabouni, S., and Nilsson, O. 2017. WUSCHEL-RELATED HOMEOBOX4 (WOX4)-like genes regulate cambial cell division activity and secondary growth in *Populus* trees. New Phytol 215:642–657.

Lakehal, A., Chaabouni, S., Cavel, E., Le Hir, R., Ranjan, A., Raneshan, Z., Novak, O., Pacurar, D.I., Perrone, I., Jobert, F., et al. 2019. A Molecular Framework for the Control of Adventitious Rooting by TIR1/AFB2-Aux/IAA-Dependent Auxin Signaling in Arabidopsis. Mol Plant 12:1499–1514.

Lakehal, A., Dob, A., Rahneshan, Z., Novak, O., Escamez, S., Alallaq, S., Strnad, M., Tuominen, H., and Bellini, C. 2020a. ETHYLENE RESPONSE FACTOR 115 integrates jasmonate and cytokinin signaling machineries to repress adventitious rooting in Arabidopsis. New Phytol 228:1611–1626.

Lakehal, A., Ranjan, A., and Bellini, C. 2020b. Multiple Roles of Jasmonates in Shaping Rhizotaxis: Emerging Integrators. Methods Mol Biol 2085:3–22.

Legue, V., Rigal, A., and Bhalerao, R.P. 2014. Adventitious root formation in tree species: involvement of transcription factors. Physiol Plant 151:192–198.

Li, J., Zhang, J., Jia, H., Liu, B., Sun, P., Hu, J., Wang, L., and Lu, M. 2018. The WUSCHEL-related homeobox 5a (PtoWOX5a) is involved in adventitious root development in poplar. Tree Physiol 38:139–153.

Li, S.W., Leng, Y., and Shi, R.F. 2017. Transcriptomic profiling provides molecular insights into hydrogen peroxide-induced adventitious rooting in mung bean seedlings. BMC Genomics 18:188.

Liu, B., Wang, L., Zhang, J., Li, J., Zheng, H., Chen, J., and Lu, M. 2014a. WUSCHEL-related Homeobox genes in *Populus* tomentosa: diversified expression patterns and a functional similarity in adventitious root formation. BMC Genomics 15:296.

Liu, J., Sheng, L., Xu, Y., Li, J., Yang, Z., Huang, H., and Xu, L. 2014b. WOX11 and 12 are involved in the first-step cell fate transition during de novo root organogenesis in Arabidopsis. Plant Cell 26:1081–1093.

Liu, S., Yang, C., Wu, L., Cai, H., Li, H., and Xu, M. 2020. The peu-miR160a-PeARF17.1/PeARF17.2 module participates in the adventitious root development of poplar. Plant Biotechnol J 18:457–469.

Love, M.I., Huber, W., and Anders, S. 2014. Moderated estimation of fold change and dispersion for RNA-seq data with DESeq2. Genome Biol 15:550.

Mauriat, M., Petterle, A., Bellini, C., and Moritz, T. 2014. Gibberellins inhibit adventitious rooting in hybrid aspen and Arabidopsis by affecting auxin transport. Plant J 78:372–384.

Merret, R., Moulia, B., Hummel, I., Cohen, D., Dreyer, E., and Bogeat-Triboulot, M.B. 2010. Monitoring the regulation of gene expression in a growing organ using a fluid mechanics formalism. BMC Biol 8:18.

Monteuuis, O., Doulbeau, S., and Verdeil, J.L. 2008. DNA methylation in different origin clonal offspring from a mature Sequoiadendron giganteum genotype. Trees-Struct Funct 22:779–784.

Nag, S., Paul, A., and Choudhuri, M.A. 2013. Changes in peroxidase activity during adventitious root formation at the base of mung bean cuttings. Int. J. Sci. Technol. Res 2:171–177.

Nilsson, O., T, A., Sitbon, F., Anthony Little, C.H., Chalupa, V., Sandberg, G., and Olsson, O. 1992. Spatial pattern of cauliflower mosaic virus 35S promoter-luciferase expression in transgenic hybrid aspen trees monitored by enzymatic assay and non-destructive imaging. TransgenicResearch 1:209–220.

Ragauskas, A.J., Williams, C.K., Davison, B.H., Britovsek, G., Cairney, J., Eckert, C.A., Frederick, W.J., Jr., Hallett, J.P., Leak, D.J., Liotta, C.L., et al. 2006. The path forward for biofuels and biomaterials. Science 311:484–489.

Ramirez-Carvajal, G.A., Morse, A.M., Dervinis, C., and Davis, J.M. 2009. The cytokinin type-B response regulator PtRR13 is a negative regulator of adventitious root development in *Populus*. Plant Physiol 150:759–771.

Ribeiro, C.L., Silva, C.M., Drost, D.R., Novaes, E., Novaes, C.R., Dervinis, C., and Kirst, M. 2016. Integration of genetic, genomic and transcriptomic information identifies putative regulators of adventitious root formation in *Populus*. BMC Plant Biol 16:66.

Rigal, A., Yordanov, Y.S., Perrone, I., Karlberg, A., Tisserant, E., Bellini, C., Busov, V.B., Martin, F., Kohler, A., Bhalerao, R., et al. 2012. The AINTEGUMENTA LIKE1 homeotic transcription factor PttIL1 controls the formation of adventitious root primordia in poplar. Plant Physiol 160:1996–2006.

Schrader, J., Nilsson, J., Mellerowicz, E., Berglund, A., Nilsson, P., Hertzberg, M., and Sandberg, G. 2004. A high-resolution transcript profile across the wood-forming meristem of poplar identifies potential regulators of cambial stem cell identity. Plant Cell 16:2278–2292.

Shukla, V., Lombardi, L., Pencik, A., Novak, O., Weits, D.A., Loreti, E., Perata, P., Giuntoli, B., and Licausi, F. 2020. Jasmonate Signalling Contributes to Primary Root Inhibition Upon Oxygen Deficiency in Arabidopsis thaliana. Plants (Basel) 9.

Sorin, C., Bussell, J.D., Camus, I., Ljung, K., Kowalczyk, M., Geiss, G., McKhann, H., Garcion, C., Vaucheret, H., Sandberg, G., et al. 2005. Auxin and light control of adventitious rooting in Arabidopsis require ARGONAUTE1. Plant Cell 17:1343–1359.

Sun, P., Jia, H., Zhang, Y., Li, J., Lu, M., and Hu, J. 2019. Deciphering Genetic Architecture of Adventitious Root and Related Shoot Traits in *Populus* Using QTL Mapping and RNA-Seq Data. Int J Mol Sci 20.

Sundell, D., Street, N.R., Kumar, M., Mellerowicz, E.J., Kucukoglu, M., Johnsson, C., Kumar, V., Mannapperuma, C., Delhomme, N., Nilsson, O., et al. 2017. AspWood: High-Spatial-Resolution Transcriptome Profiles Reveal Uncharacterized Modularity of Wood Formation in *Populus tremula*. Plant Cell 29:1585–1604.

Supek, F., Bosnjak, M., Skunca, N., and Smuc, T. 2011. REVIGO summarizes and visualizes long lists of gene ontology terms. PLoS One 6:e21800.

Taeroe, A., Nord-Larsen, T., Stupak, I., and Raulund-Rasmussen, K. 2015. Allometric biomass, biomass expansion factor and wood density models for the OP42 hybrid poplar in southern Scandinavia. BioEnergy Research 8:1332–1343.

Team, R.C. 2018. R: A language and environment for statistical computing. In: Available online at https://www.R-project.org/--R Foundation for Statistical Computing, V., Austria, ed.

Trupiano, D., Yordanov, Y., Regan, S., Meilan, R., Tschaplinski, T., Scippa, G.S., and Busov, V. 2013. Identification, characterization of an AP2/ERF transcription factor that promotes adventitious, lateral root formation in *Populus*. Planta 238:271–282.

Tuskan, G.A., and Difazio, S., and Jansson, S., and Bohlmann, J., and Grigoriev, I., and Hellsten, U., and Putnam, N., and Ralph, S., and Rombauts, S., and Salamov, A., et al.2006. The genome of black cottonwood, *Populus* trichocarpa (Torr. & Gray). Science 313:1596–1604.

Velada, I., Grzebelus, D., Lousa, D., C, M.S., Santos Macedo, E., Peixe, A., Arnholdt-Schmitt, B., and H, G.C. 2018. AOX1-Subfamily Gene Members in Olea europaea cv. “Galega Vulgar”-Gene Characterization and Expression of Transcripts during IBA-Induced in Vitro Adventitious Rooting. Int J Mol Sci 19.

Wang, J.W. 2014. Regulation of flowering time by the miR156-mediated age pathway. J Exp Bot 65:4723–4730.

Wang, L.-Q., Li, Z., Wen, S.-S., Wang, J.-N., Zhao, S.-T., and Lu, M.-Z. 2020. WUSCHEL-related homeobox gene PagWOX11/12a responds to drought stress by enhancing root elongation and biomass growth in poplar. Journal of Experimental Botany 71:1503–1513.

Wei, M., Liu, Q., Wang, Z., Yang, J., Li, W., Chen, Y., Lu, H., Nie, J., Liu, B., Lv, K., et al. 2020. PuHox52-mediated hierarchical multilayered gene regulatory network promotes adventitious root formation in *Populus ussuriensis*. New Phytol 228:1369–1385.

Willmann, M.R., and Poethig, R.S. 2005. Time to grow up: the temporal role of smallRNAs in plants. Current opinion in plant biology 8:548–552.

Wuddineh, W.A., Mazarei, M., Turner, G.B., Sykes, R.W., Decker, S.R., Davis, M.F., and Stewart, C.N., Jr. 2015. Identification and Molecular Characterization of the Switchgrass AP2/ERF Transcription Factor Superfamily, and Overexpression of PvERF001 for Improvement of Biomass Characteristics for Biofuel. Front Bioeng Biotechnol 3:101.

Xu, C., Tao, Y., Fu, X., Guo, L., Xing, H., Li, C., Yang, Z., Su, H., Wang, X., Hu, J., et al. 2021. The microRNA476a-RFL module regulates adventitious root formation through a mitochondria-dependent pathway in *Populus*. New Phytol.

Xu, M., Xie, W., and Huang, M. 2015. Two WUSCHEL-related HOMEOBOX genes, PeWOX11a and PeWOX11b, are involved in adventitious root formation of poplar. Physiol Plant 155:446–456.

Yordanov, Y.S., Ma, C., Yordanova, E., Meilan, R., Strauss, S.H., and Busov, V.B. 2017. BIG LEAF is a regulator of organ size and adventitious root formation in poplar. PLoS One 12:e0180527.

Yue, J., Yang, H., Yang, S., and Wang, J. 2020. TDIF regulates auxin accumulation and modulates auxin sensitivity to enhance both adventitious root and lateral root formation in poplar trees. Tree Physiol 40:1534–1547.

Zhang, B., Tong, C., Yin, T., Zhang, X., Zhuge, Q., Huang, M., Wang, M., and Wu, R. 2009. Detection of quantitative trait loci influencing growth trajectories of adventitious roots in *Populus* using functional mapping. Tree genetics & genomes 5:539–552.

Zhang, Y., Xiao, Z., Zhan, C., Liu, M., Xia, W., and Wang, N. 2019. Comprehensive analysis of dynamic gene expression and investigation of the roles of hydrogen peroxide during adventitious rooting in poplar. BMC Plant Biol 19:99.

Zhang, Y., Yang, X., Cao, P., Xiao, Z., Zhan, C., Liu, M., Nvsvrot, T., and Wang, N. 2020. The bZIP53-IAA4 module inhibits adventitious root development in *Populus*. J Exp Bot 71:3485–3498.

