## Supporting Information for "Molecular basis of differential adventitious rooting competence in poplar genotypes"

**The following Supporting Information is available for this article:**

**Methods S1** Plant growth conditions and rooting assays

**Methods S2** Histological analysis of stem cuttings *in vitro*

**Methods S3** Tissue preparation before laser capture microdissection

**Methods S4** Pre-processing of RNA-Seq data

**Methods S5** Generation of plasmid constructs and transformation of hybrid aspen

**Methods S6** Quantitative Real-Time PCR analysis

**Fig. S1** Conditions for adventitious rooting assays from *in vitro* plants and greenhouse-grown plants

**Fig. S2:** Workflow for laser capture microdissection (LCMS) of cambium tissues from stem **cuttings**

**Fig. S3**: Quality assessment of the RNAseq data in the different biological replicates

**Fig. S4**: Populus Arabidopsis orthologues of *ARF6*, *ARF8* and *ARF17* and their expression pattern in wood-forming tissues

**Fig.S5:** Heat map showing the average expression of genes encoding ROS scavenging proteins in the cambium of T89 and OP42 genotypes

**Fig.S6:** Heat map showing the average expression of *PtrARF* genes in the cambium of T89 and OP42 genotypes

**Fig. S7: Over-expression** of *PtAF6.4* and PtARF8.2 under the 35S promoter

**Table S1** Primer list used in the present study.

**Dataset S1 (**excel document):

- **Sheet 1** Library-size-normalised variance-stabilised data set
- **Sheet 2** Expression values for the 17,997 expressed genes

**Dataset S2:** Differentially expressed genes

**Sheet 1** The differentially expressed genes (DEG) up- and down-regulated and their annotation.

**Sheet** 2 Total number of DEG in OP42 when compared at time T1 and T0.

**Sheet 3** The DEG up-regulated in OP42 at time T1 compared to time T0.

**Sheet 4** The DEG down-regulated in OP42 at time T1 compared to time T0.

**Sheet 5** Total number of DEG in T89 when compared at time T1 and T0.

**Sheet 6** The DEG up-regulated in T89 at time T1 compared to time T0.

**Sheet 7** The DEG down-regulated in T89 at time T1 compared to time T0.

**Sheet 8** The total number of DEG in OP42 and T89 at time T0.

**Sheet 9** The number of DEG up-regulated in T89 compared to OP42 at timeT0.

**Sheet 10** The number of DEG down-regulated in T89 compared to OP42 at timeT0.

**Sheet 11** Total number of DEG between T89 and OP48 at time T1.

**Sheet 12** The number DEG up-regulated in T89 compared to OP42 at time T1.

**Sheet 13** The number of DEG down-regulated in T89 compared to OP42 at time T1.

**Dataset S3**

**Sheet 1** Vascular tissue expressed genes set.

**Sheet 2** Differentially expressed Transcription Factors.

**Sheet 3** Differentially expressed ROS scavenging proteins

**Sheet 4** Gene Ontology of up-regulated DEGs.

**Sheet 5** Gene Ontology of down-regulated DEGs.

**Sheet 6** Mean of expression values used for the heat maps

**Methods S1** Plant growth conditions and rooting assays

The hybrid aspen (*Populus tremula L.* × *Populus tremuloides Michx*), clone T89 was micropropagated *in vitro* for four weeks, on sterile half-strength Murashige and Skoog (1/2 MS) medium (pH 5.6) (Duchefa, https://www.duchefa-biochemie.com/) as described in (Karlberg *et al.*, 2011) in plastic jars at 25 °C ± 1 °C under a 18:6 h light/dark cycle provided by white fluorescent artificial light with 50 µmol/m$^2$/s light intensity in a growth chamber. For *in vitro* rooting assays, 3 cm cuttings with four to five leaves in the case of T89, and two to three leaves in the case of *P.trichocarpa*× *P.*maximowiczii clone OP42 plantlets, were collected and transferred in smaller rectangular jars containing fresh sterile medium.

For the rooting assay in hydroponic conditions, four-week-old *in vitro* T89 and OP42 plantlets were transferred to soil and kept in the greenhouse for three months (16 h light, 21°C; 8 h dark 18 °C). 20 cm lengths of stem cuttings were taken from the third internode below the shoot apex. After removal of all leaves and buds except for the higher axillary bud (Figs S1c, e), the cuttings were transferred to hydroponic conditions in the greenhouse. The nutrient solution was composed of a modified Hoagland solution as described in Plett *et al.* (2011). Photos of the AR were taken using a Canon EOS 350 digital camera and Discovery V.8 stereomicroscope fitted with a Zeiss camera.

**Methods S2** Histological analysis of stem cuttings *in vitro*

For histological analysis of stems, 5 mm stem fragments were taken at the base of cuttings four or five days after cutting. Samples were vacuum infiltrated with a fixation medium (10 ml of 37% formaldehyde, 5ml of 5% acetic acid, 50 ml of 100% ethanol and 35 ml of $H_2O$) for 20 seconds and left for 24 h at room temperature. The samples were then washed in 70% ethanol for 10 minutes and transferred into fresh 70% EtOH until required for use. Samples were then gradually dehydrated in an ethanol series (80%, 90%, 96% for 2 h each and 100% overnight at room temperature). The 100% EtOH was gradually replaced by HistoChoice tissue fixative (VWR Life, https://us.vwr.com/) in three steps of 1:3, 1:1, 3:1 (EtOH: HistoChoice ratio), then with pure HistoChoice twice in 1 h. The HistoChoice fixative was gradually replaced with Paraplast Plus for tissue embedding (Sigma-Aldrich, https://www.sigmaaldrich.com/), over six days.

**Methods S3** Tissue preparation before laser capture microdissection

*Sampling, fixation and cryoprotection steps*

The basal 5 mm stem pieces of T89 and OP42 cuttings were harvested immediately after excision from greenhouse-grown plants (Time T0) and after 24 h of hydroponic culture (Time T1) (Figs S2a-c). Three biological replicates of tissue samples were collected at each time point (T0 and T1) from both OP42 and T89 (12 samples in total = 3 biological replicates x 2 genotypes x 2 time points). Immediately after the sampling, stem pieces were split in half longitudinally and subjected to fixation and cryoprotection steps before the laser microdissection. We used the protocol described at https://schnablelab.plantgenomics.iastate.edu/resources/protocols/, slightly modified as follows: samples were soaked in cold Ethanol-Acetic Acid (EAA) Farmer's fixative solution, containing 75% (v/v) ethanol and 25% (v/v) acetic acid, and vacuum infiltrated on ice at 400 mm Hg for 20 minutes. After 1 h incubation at 4 °C, another step of vacuum infiltration with fresh Farmer`s solution was performed (400 mm Hg for 20 min). Samples were then kept at 4 °C overnight. The following day, the fixative solution was removed and the samples transferred in a 10% sucrose solution prepared with 1X Phosphate Buffered Saline (PBS) (137 mM NaCl, 8 mM Na2PO4, 2.68 mM KCl, 1.47 mM KH2PO4), vacuum infiltrated on ice at 400 mm Hg for 15 min. Samples were left incubating for 1 h at 4 °C, then vacuum infiltrated with a 15% sucrose solution (400 mm Hg for 15 min). Samples were then incubated overnight at 4 °C; then frozen in liquid nitrogen and stored at -80 °C until cryosectioning.

*Cryosectioning*

The day before cryosectioning, membrane slides for laser microdissection (FrameSlide PET, Zeiss; https://www.fishersci.co.uk/) were treated with RNaseZap (Sigma, https://www.sigmaaldrich.com/), rinsed twice with diethylpyrocarbonate (DEPC) water and dried for 2 h at 37 °C. Immediately before sectioning, slides were further treated with UV light for 30 min to improve sections adhesion. Tweezers and a cryostat knife were sterilised at 180 °C for 4 h. The chamber temperature of the cryostat (Leica CM1850) was set at -25 °C. The instruments including tweezers, knives, and Polyethylene Teraphthalate (PET)-membrane coated slides were transferred into the chamber 20 min before sectioning. Samples were transferred from -80°C freezer to the cryostat in liquid nitrogen. They were fixed with Tissue-Tek® Optimal Cutting Temperature (O.C.T.) compound onto a specimen stage directly in the cryochamber. To avoid embedding and the presence of O.C.T. compound on membrane slides, stem segments were mounted to allow cambium collection from tangential cryosections (Fig. S2d). Sections of 25 µm were transferred with tweezers onto membrane slides then moved in a Petri dish at room temperature. Sections were then treated with 70% ethanol for 5 min at room temperature, followed by 95% ethanol for 2 min on ice, and 100% ethanol for 2 min on ice. In these dehydration steps ethanol was applied and removed directly onto the membrane slide chamber with a sterile plastic Pasteur pipette, being careful not to damage the membrane. After ethanol removal, sections were air-dried for 5 min before being cut at the microdissector.

**Methods S4** Pre-processing of RNA-Seq data

The data pre-processing was performed as described in: http://www.epigenesys.eu/en/protocols/bio-informatics/1283-guidelines-for-rna-seq-data.
Briefly, the quality of the raw sequence data was assessed using FastQC (http://www.bioinformatics.babraham.ac.uk/projects/fastqc/).
Residual ribosomal RNA (rRNA) contamination was assessed and filtered using SortMeRNA (v2.1; Kopylova *et al*., 2012; settings --log –paired in --fastx--sam --num_alignments 1) using the rRNA sequences provided with SortMeRNA (rfam-5s-database-id98.fasta, rfam-5.8s-database-id98.fasta, silva-arc-16s-database-id95.fasta, silva-bac-16s-database-id85.fasta, silva-euk-18s-database-id95.fasta, silva-arc-23s-database-id98.fasta, silva-bac-23s-database-id98.fasta and silva-euk-28s-database-id98.fasta). Data were then filtered to remove adapters and trimmed for quality using Trimmomatic (v0.32; Bolger *et al*., 2014; settings TruSeq3-PE-2.fa:2:30:10 LEADING:3 SLIDINGWINDOW:5:20 MINLEN:50). After both filtering steps, FastQC was run again to ensure that no technical artefacts were introduced. Filtered reads were aligned to v3.0 of the *P. trichocarpa* genome (Phytozome) using STAR (v2.5.2b; Dobin *et al*., 2013; non default settings: --outSAMstrandField intronMotif--readFilesCommand zcat--outSAMmapqUnique 254 --quantMode TranscriptomeSAM --outFilterMultimapNmax 100 --outReadsUnmapped Fastx --chimSegmentMin1--outSAMtype BAM SortedByCoordinate --outWigType bedGraph --alignIntronMax 11000). The annotations obtained from the *P. trichocarpa* v3.0 GFF file were flattened to generate 'synthetic' gene models. This synthetic transcript GFF file and the STAR read alignments were used as input to the HTSeq (Anders *et al*., 2015) htseq-count python utility to calculate exon-based read count values. The htseq-count utility takes only uniquely mapping reads into account.

**Methods S5** Generation of plasmid constructs and transformation of hybrid aspen

To amplify the candidate genes, cDNA was synthesised (SuperScript II Reverse Transcriptase, Invitrogen) starting from total RNA extracted from hybrid aspen T89 (*P. tremula* x *P. tremoloides*) leaves using Spectrum™ Plant Total RNA Kit (Sigma-Aldrich) followed by DNAse treatment (TURBO DNA-free Kit, Ambion). As it is not possible to distinguish the *P. tremula* sequence from that of *P. tremuloides*, the genes are referred to as *PttARF6.4*, *PttARF8.2*, *PttARF17.2* and *PttMYC2.1* and the corresponding primers used for amplification of the coding sequences are listed in Table S1.

The amplified cDNA of *PttARF6.4, PttARF8.2* and *PttMYC2.1* were cloned independently into the pENTR/D-TOPO donor vector (https://www.fishersci.se/se/en/home.html) and transferred into the pK2GWF7 plant transformation vector. *PttARF6.4* and *PttARF8.2* coding sequences were also cloned in the pK2GWFS7 vector in which the CaMV35S promoter had been replaced by a 2-kb promoter fragment from the *PttHB3a* gene for specific expression in the cambium (Schrader *et al*., 2004). To down-regulate the *ARF*s genes we generated RNAi constructs with 578 bp, 624 bp and 480 bp fragments from *PttARF6.4*, *PttARF8.2* and *PttARF17.2*, respectively. These fragments were amplified using primers listed in Table S1 and T89 cDNA as a template. Due to high coding nucleotide sequence similarity, RNAi constructs targeting both *Ptt*A*RF6.3* and *PttARF6.4* paralogues, *PttARF8.1* and *PttARF8.2* paralogues or *PttARF17.1* and *PttARF17.2* paralogues were generated. The amplified fragments were cloned into pENTR/D-TOPO (Invitrogen) and then transferred into the plant transformation vectors pK7GWIWG2.

All the different constructs were transformed independently into *Agrobacterium tumefaciens* GV3101 pmp90RK, which were used to transform the hybrid aspen T89. In total, 14 independently transformed lines for each construct were generated. The relative expression levels of *PttARF6.1/2*, *PttARF6.3/4, Ptt*ARF*17.1/2* and *PttARF17.1/2* in the respective transgenic lines were further quantified by qPCR. Two independent RNAi lines for each construct were selected and analysed for their adventitious rooting ability.

**Methods S6** Quantitative Real-Time PCR analysis

Total RNA was extracted from the base of five pieces of 5 mm stem cuttings of T89 and transgenic lines that were collected at the time of the adventitious rooting assay (3 biological replicates for each line, each biological replicate formed by stem pieces collected from 3 different plants). Total

RNA was extracted using the Spectrum™ Plant Total RNA Kit (Sigma-Aldrich). A total 10 μg of RNA samples was treated with TURBO DNA-free Kit (Ambion) to remove contaminating DNA from RNA preparations, and to remove the DNAse from the samples. cDNA was synthesised using SuperScript® III Reverse Transcriptase Kit (Invitrogen) following the DNase treatment. Quantitative real-time PCR analyses were carried out with a Roche LightCycler 480 II instrument, and expression values were calculated relative to the reference gene expression values, by using the Δ-ct-method as previously described by (Gutierrez *et al.*, 2008).

**References:**

**Anders, S., Pyl, P.T., and Huber, W. 2015**. HTSeq--a Python framework to work with high-throughput sequencing data. *Bioinformatics* **31**:166-169.

**Bolger, A.M., Lohse, M., and Usadel, B. 2014**. Trimmomatic: a flexible trimmer for Illumina sequence data. *Bioinformatics* **30**:2114-2120

**Gutierrez, L., Mauriat, M., Guenin, S., Pelloux, J., Lefebvre, J.F., Louvet, R., Rusterucci, C., Moritz, T., Guerineau, F., Bellini, C., and Van Wuytswinkel, O. 2008**. The lack of a systematic validation of reference genes: a serious pitfall undervalued in reverse transcription-polymerase chain reaction (RT-PCR) analysis in plants. *Plant Biotechnol J* **6**:609-618

**Karlberg, A., Bako, L., and Bhalerao, R.P. 2011**. Short day-mediated cessation of growth requires the down regulation of AINTEGUMENTALIKE1 transcription factor in hybrid aspen. *PLoS Genet* **7**:e1002361.

**Kopylova, E., Noe, L., and Touzet, H. 2012**. SortMeRNA: fast and accurate filtering of ribosomal RNAs in metatranscriptomic data. *Bioinformatics* **28**:3211-3217.

**Plett, J.M., Montanini, B., Kohler, A., Ottonello, S., and Martin, F. 2011**.Tapping genomics to unravel ectomycorrhizal symbiosis. *Methods Mol Biol.* 722:249-81

**Schrader, J., Nilsson, J., Mellerowicz, E., Berglund, A., Nilsson, P., Hertzberg, M., and Sandberg, G. 2004**. A high-resolution transcript profile across the wood-forming meristem of poplar identifies potential regulators of cambial stem cell identity. *Plant Cell* **16**:2278-2292.

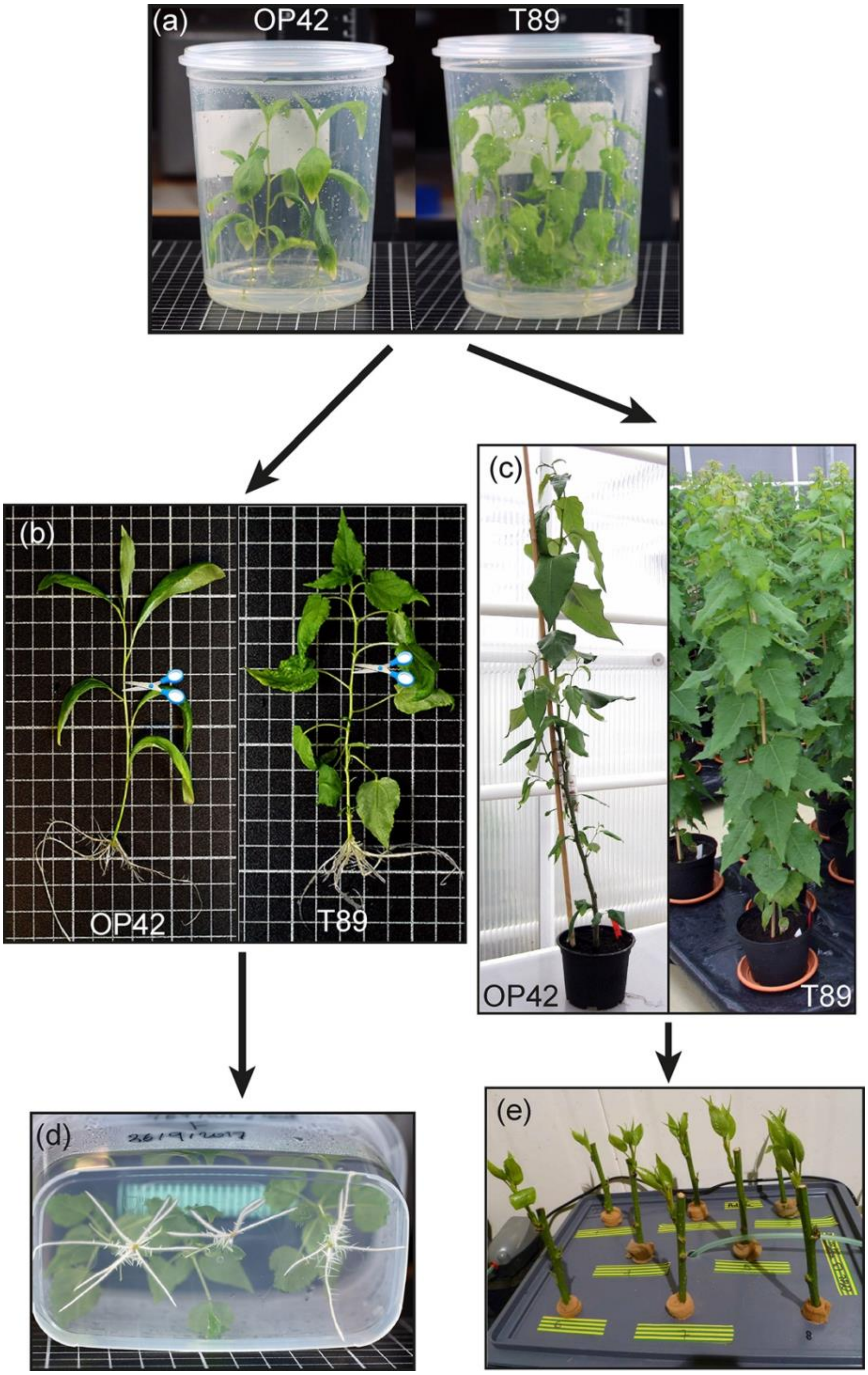


**Fig. S1: Conditions for adventitious rooting assays from *in vitro* plants and greenhouse-grown plants**

(a) OP42 and T89 plants are propagated under *in vitro* conditions for four weeks.

(b, d) Cuttings comprising the shoot apex and the three first internodes starting from the shoot apex were excised from 4-week-old plants (b) and transferred to fresh ½ MS medium in smaller rectangular jars (d). The number of AR was monitored, starting 5 days after being cut, when the first macroscopic events could be observed at the base of the cuttings, until 14 days after cutting as in (d).

(c) Four-week-old *in vitro* OP42 and T89 plants were transferred into pots containing soil and left to grow for three months in the greenhouse. (e) Approximately 20 cm stem cuttings with a 1 cm stem diameter were excised from the three-month-old plants and transferred in hydroponic conditions.

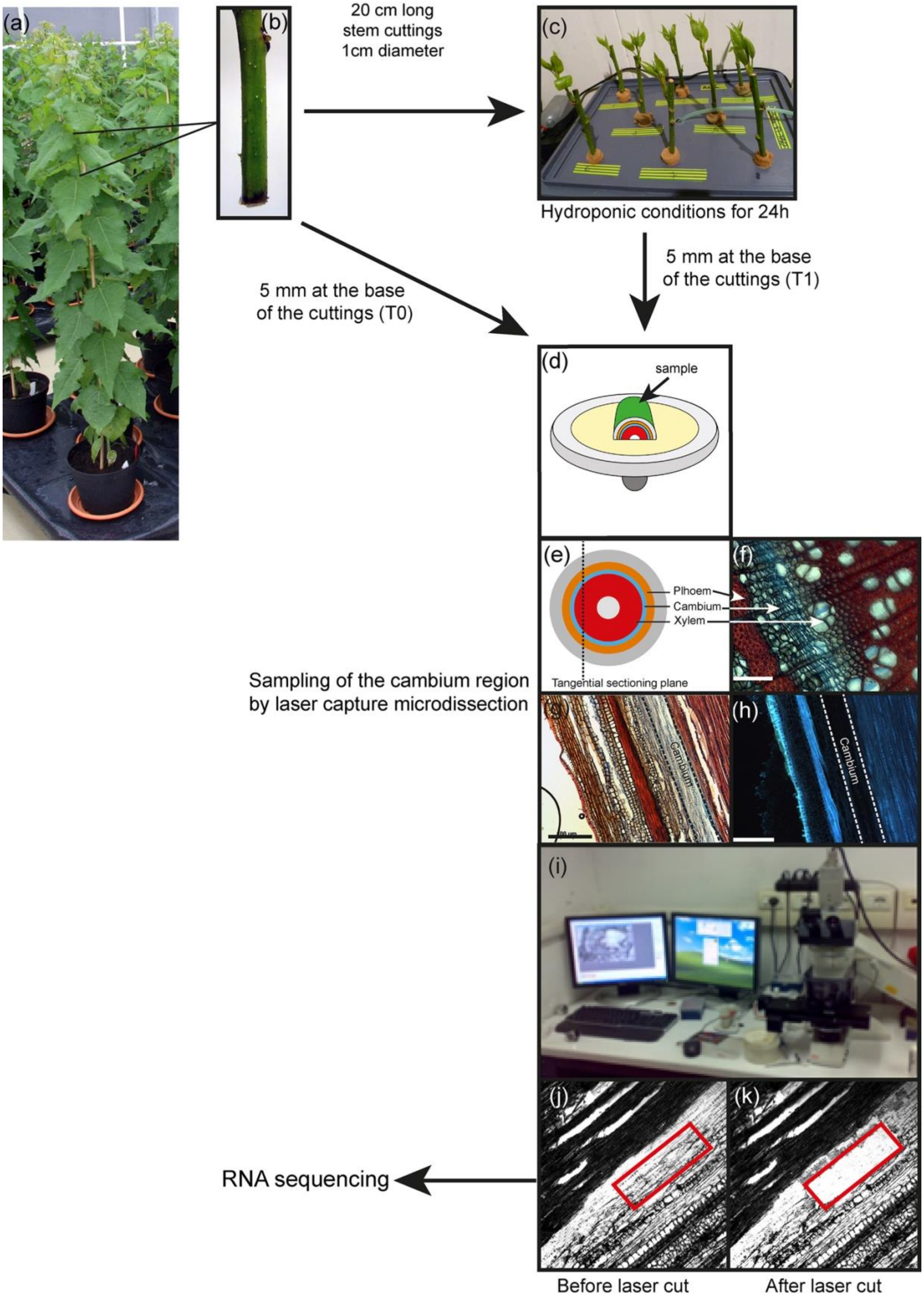

(a)
(b)
20 cm long
stem cuttings
1cm diameter
(c)
Hydroponic conditions for 24h
5 mm at the base
of the cuttings (T1)
5 mm at the base
of the cuttings (T0)
(d)
sample
(e)
(f)
Plhoem
Cambium
Xylem
Tangential sectioning plane
Sampling of the cambium region
by laser capture microdissection
(g)
Cambium
(h)
Cambium
(i)
(j)
(k)
RNA sequencing
Before laser cut
After laser cut

**Fig. S2: Workflow for laser capture microdissection (LCMS) of cambium tissues from stem cuttings**

(a) T89 and OP42 plants were grown in the greenhouse for 3 months.

(b) 20 cm lengths of stem cuttings were taken as for the hydroponic assay (Supplementary Figure 1) and 5 mm long pieces were cut at their base, flash frozen in liquid nitrogen and used for cambium tissue sampling at time T0.

(c) A second set of stem cuttings were kept in hydroponic conditions for 24 h (T1) and 5 mm long stem pieces were cut at the base of the cuttings, flash frozen in liquid nitrogen and used for cambium tissue sampling at time T1.

(d) 5 mm stem pieces were split in half longitudinally.

(e) Schematics of the anatomy of a stem; (f) cross-section of a stem cutting; (g and h) Longitudinal cryosection of the base of a stem cutting observed under the microscope of the micromanipulator with white light (g) or UV light which allowed us to identify more precisely the cambium region which did not show any fluorescence (h). (i) computerised system for LCMS. Cambium region before (j) and after (k) laser microdissection.

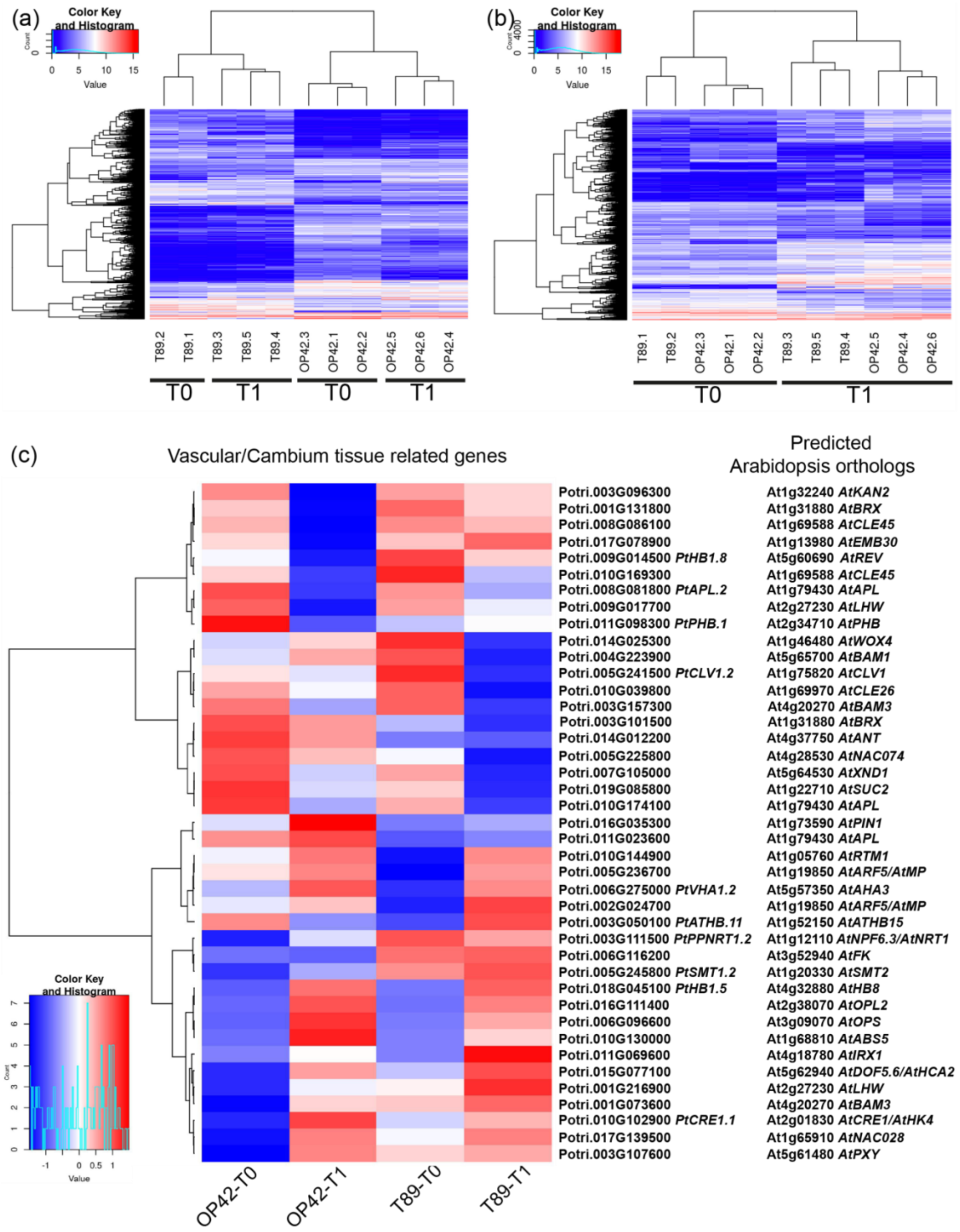


**Fig. S3: Quality assessment of the RNAseq data in the different biological replicates**

(a) The dendrogram of samples (top) was divided into two parts based on the correlation between a genotype's gene expression and then labelled (bottom), respectively. (b) The dendrogram of samples (top) was divided into two parts based on the correlation between time and a treatment's gene expression and then labeled (bottom), respectively. (c) The heat map was generated based on genotypes (T89 and OP42) and time after cutting. T0 immediately after cutting, and T1 24 h after cutting and being transferred to hydroponic conditions. Heatmaps of DE genes (DE cut-offs of FDR $\leq 0.01$ and $|LFC| \geq 0.5$), were generated using the function heatmap.2 from the gplots R library. The genes, which were expressed in either one or two biological replicates, but which expression was significantly different between T89 and OP42, were also mapped with the variance stabilising transform (VST) data set. The gene expression mean values used for the heat map are listed in Supplementary data set 3, sheet 6.

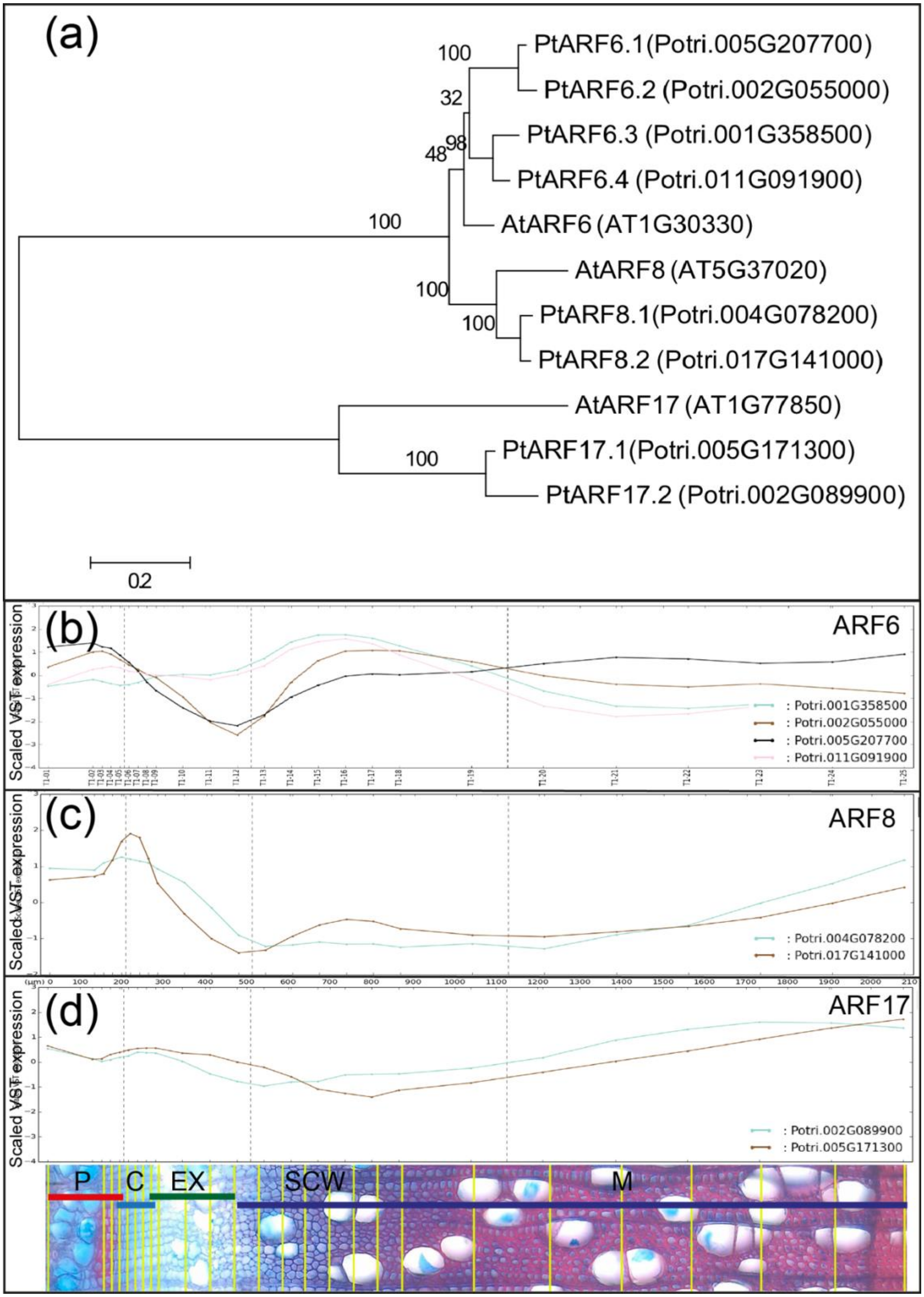

(a)
PtARF6.1(Potri.005G207700)
PtARF6.2 (Potri.002G055000)
PtARF6.3 (Potri.001G358500)
PtARF6.4 (Potri.011G091900)
AtARF6 (AT1G30330)
AtARF8 (AT5G37020)
PtARF8.1(Potri.004G078200)
PtARF8.2 (Potri.017G141000)
AtARF17 (AT1G77850)
PtARF17.1(Potri.005G171300)
PtARF17.2 (Potri.002G089900)
100
32
98
48
100
100
100
100
02
(b)
ARF6
Scaled VST expression
: Potri.001G358500
: Potri.002G055000
: Potri.005G207700
: Potri.011G091900
(c)
ARF8
Scaled VST expression
: Potri.004G078200
: Potri.017G141000
(d)
ARF17
Scaled VST expression
: Potri.002G089900
: Potri.005G171300
P
C
EX
SCW
M

**Fig. S4**: **Populus Arabidopsis orthologues of ARF6, ARF8 and ARF17 and their expression pattern in wood-forming tissues**

(a) Phylogenetic relationship between *P. trichocarpa* and *Arabidopsis thaliana* ARF6, ARF8 and ARF17 proteins. Protein sequences were aligned with ClustalW and the phylogenetic analysis was performed in Mega 8 using the Neighbour-Joining method with a bootstrap test (1000 replicates). (b-d) Expression patterns of *ARF6*, *ARF8* and *ARF17* genes in the wood-forming regions of aspen trees (http://aspwood.popgenie.org). The y-axis shows the variance-scaled expression. The x-axis shows tangential samples over the wood-developing tissues with four zones indicated: P = phloem; C + EX = cambium and expansion zones; SCW = secondary cell wall deposition zone; M = maturation zone (Sundell *et al.*, 2017). The corresponding Potri. identifications are *PtARF6.1*, Potri.005G207700; *PtARF6.2*, Potri.002G055000; *PtARF6.3*, Potri. 001G358500; *PtARF6.4*, Potri.011G091900; *PtARF8.1*, Potri.004G078200; *PtARF8.2*, Potri.017G141000; *PtARF17.1*, Potri.005G171300; *PtARF17.2*, Potri.002G089900.

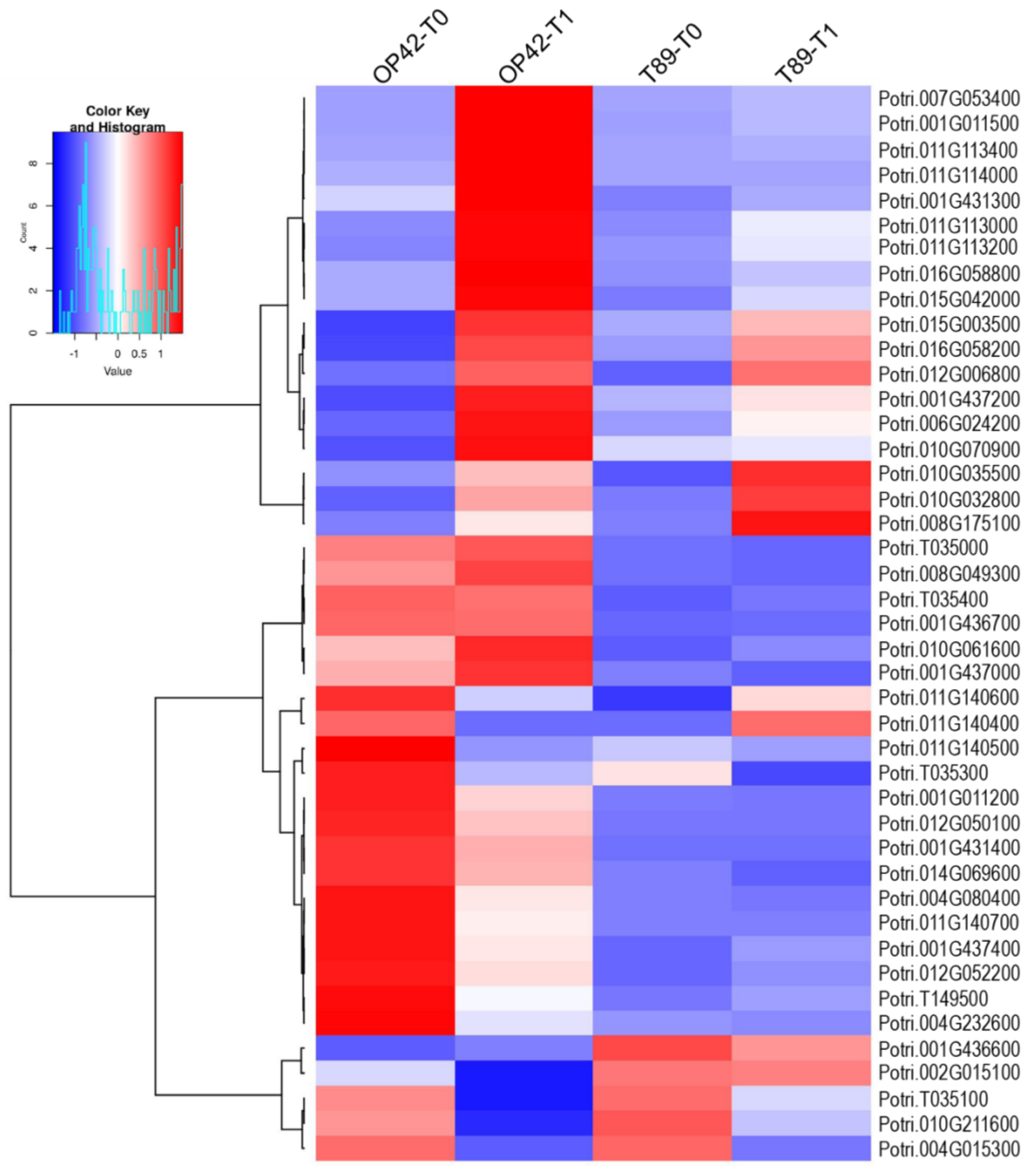


**Fig. S5: Heat map showing the average expression of genes encoding ROS scavenging proteins in the cambium of T89 and OP42 genotypes**

The heat map was generated based on genotypes (T89 and OP42) and time after cutting. T0 immediately after cutting and T1 24 h after cutting and being transferred to hydroponic conditions. (Supplementary data sets 2 and 3, sheets 3 and 4). The heat map was generated based on genotypes (T89 and OP42) and time after cutting. T0 immediately after cutting and T1 24 h after cutting and being transferred to hydroponic conditions. Heat maps of DE genes (DE cut-offs of FDR $\leq$ 0.01 and $|LFC| \geq 0.5$), were generated using the function heatmap.2 from the gplots R library. The genes, which were expressed in either one or two biological replicates, but which expression was significantly different between T89 and OP42, were also mapped with the variance stabilising transform (VST) data set. The gene expression mean values used for the heat map are listed in Supplementary data set 3, sheet 6.

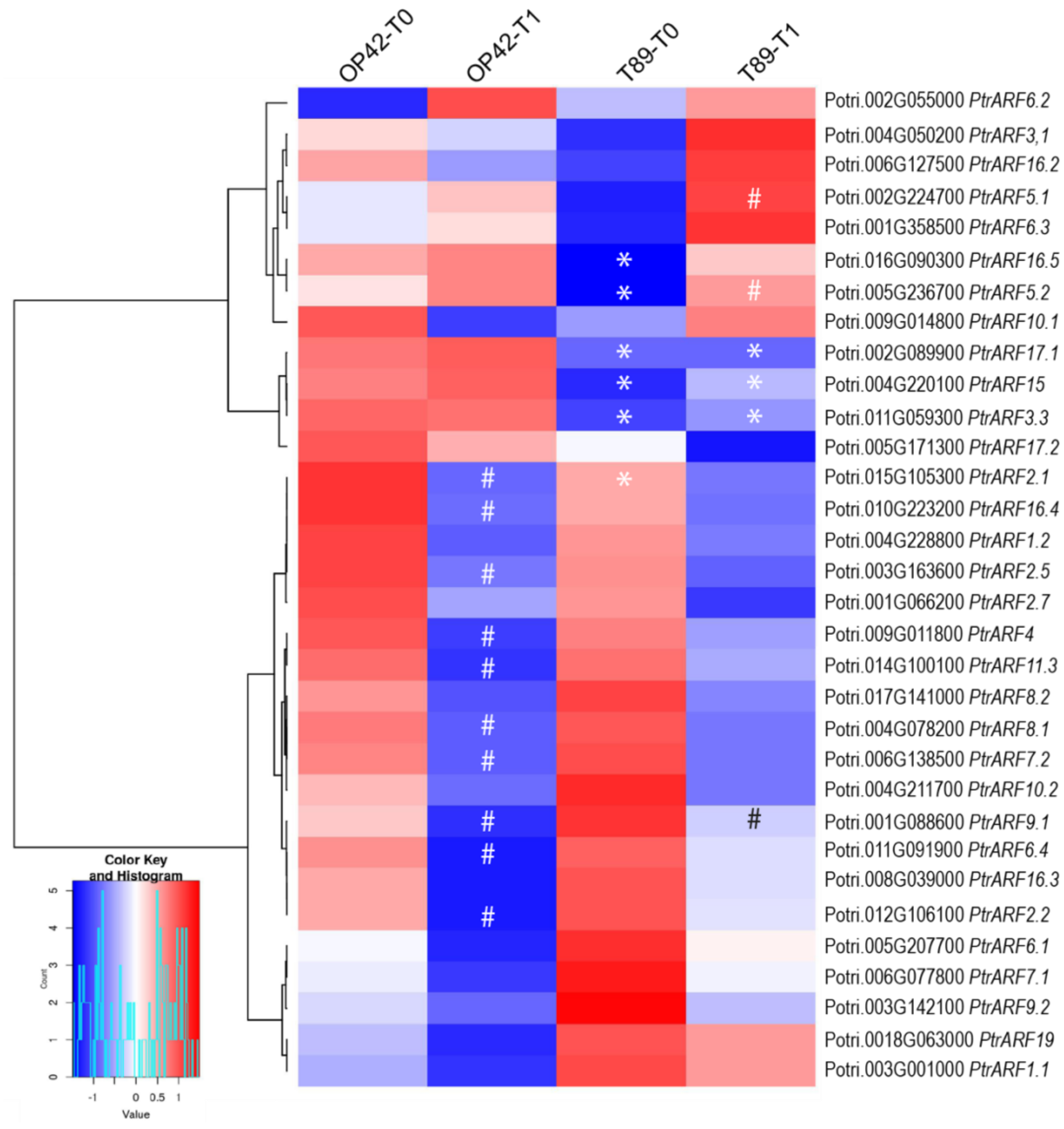


**Fig. S6: Heat map showing the average expression of *PtARF* genes in the cambium of T89 and OP42 genotypes**

The heat map was generated based on genotypes (T89 and OP42) and time after cutting. T0 immediately after cutting and T1 24 h after cutting and being transferred to hydroponic conditions. The asterisks indicate that the expression in T89 compared to OP42 at T0 or T1 is significantly different. The dashes indicate that the expression at T1 compared to T0 in either T89 or Op42 is significantly different (Supplementary data sets 2 and 3). The gene expression mean values used for the heat map are listed in Supplementary data set 3, sheet 6.

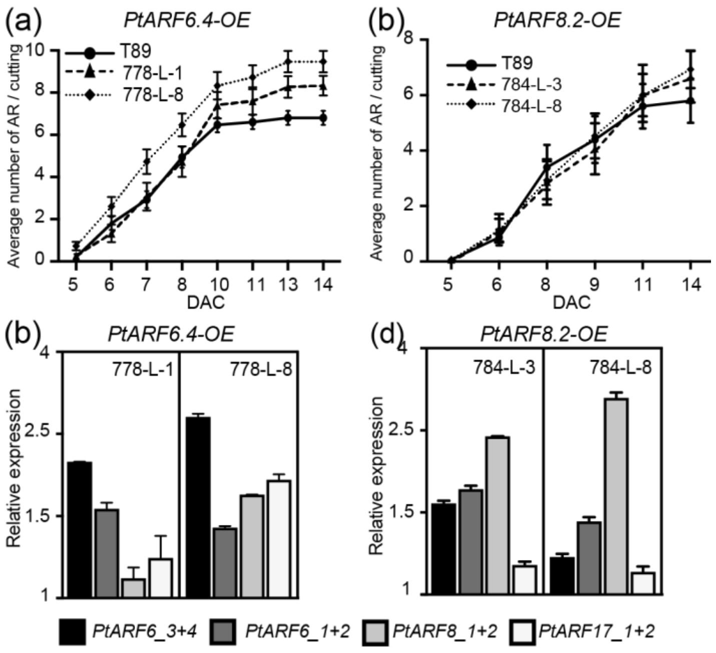


**Fig. S7: Over-expression of *PtAF6.4* and PtARF8.2 under the 35S promoter**

(a-b) Average number of AR on cuttings of transgenic plants expressing *p35S:PtARF6.4* (a) and *p35S*:*PtARF8.2* (b) Rooting assays were performed as described in Materials and Methods. Two independent transgenic lines were compared to the control T89. AR number was scored every day starting day 5 after being cut until 14 days after cutting (DAC). For each line 15 cuttings were analysed. Data are means ± SE, n = 15, corresponding to two independent lines per construct.

(c-d) The *PtARF6.1/2*, *PtARF6.3/4*, *PtARF8.1/2*, *PtARF17.1/2* un-cleaved transcript abundance was quantified in stem cutting fragments of *p35S:PtARF6.4* and *p35S*:*PtARF8.2* over-expressing lines and the control line T89. Gene expression values are relative to the reference gene and calibrated towards the expression in the control line T89, for which the value is set to 1. Error bars indicate SE obtained from three independent biological replicates. A one-way analysis of variance combined with the Dunnett's comparison post-test indicated that the values marked with an asterisk were significantly different from T89 values ($P < 0.05$; $n = 3$).

**Table S1** Primer list used in the present study.

| | | Forward Primer | Reverse Primer |
|---|---|---|---|
| | **Cloning Primers** | | |
| Potri.011G091900 | PtARF6.4 | CACCATGAGGCACTCTTCGGCTTC | TTAAATTTCTCGGCAGTCCAAAGAC |
| Potri.017G141000 | PtARF8.2 | CACCATGAAGCTTTCAACATCAGG | TCATCCTTTGACAGCATTTGGGCC |
| Potri.001G358500/ Potri.011G091900 | PtARF6.3/4 RNAi | CACCACTGCTGCGTTTCAGGAGAT | ATGAGATGTTTCGTCCTGGG |
| Potri.004G078200/ Potri.017G141000 | PtARF8.1/2RNAi | CACCCAAATTTCAACAGAAAGCTTGC | GTAGATTGACCAGCTCTGGAGA |
| Potri.005G171300/ Potri.002G089900 | PtARF17.1/2RNAi | CACCAACGGTGGTGGTTTCTCCGTC | ACCGCCACCAGCAATCTGCT |
| Potri.003G092200 | PtMYC2 | CACCATGACTGATTACCGTCTA | CTATCGGGCATCACCAACTTTTGT |
| | **qPCR Primers** | | |
| Potri.001G358500/ Potri.011G091900 | PtARF6.3/4 | GAGTTGCGAAGTGAGCTTGC | TTACAAATTCCGGCCAGGGG |
| Potri.005G207700/ Potri.002G055000 | PtARF6.1/2 | ATGATGAGCTTCGCAGTGAGC | AGGATCATCACCAAGGAGAAGC |
| Potri.004G078200/ Potri.017G141000 | PtARF8.1/2 | GGACATATCCCGGTTCAGCA | ACTCCCAGGGATCATCTCCAA |
| Potri.005G171300/ Potri.002G089900 | PtARF17.1/2 | CCCAATGAAGAAATTGAGATATCC | GAATGTGGAAAAAGGATCTTGC |
| Potri.003G092200 | PtMYC2.1 | CTACGAGCTGTGGTTCCTAATGTAT | ATTTGACATCTTAAGCTCCTGATTG |
| Potri.001G418500 | PtUBQ | GTTGATTTTTGCTGGGAAGC | GATCTTGGCCTTCACGTTGT |
